# Dietary isoleucine content defines the metabolic and molecular response to a Western diet

**DOI:** 10.1101/2024.05.30.596340

**Authors:** Michaela E. Trautman, Cara L. Green, Michael R. MacArthur, Krittisak Chaiyakul, Yasmine H. Alam, Chung-Yang Yeh, Reji Babygirija, Isabella James, Michael Gilpin, Esther Zelenovskiy, Madelyn Green, Ryan N. Marshall, Michelle M. Sonsalla, Victoria Flores, Judith A. Simcox, Irene M. Ong, Kristen C. Malecki, Cholsoon Jang, Dudley W. Lamming

## Abstract

The amino acid composition of the diet has recently emerged as a critical regulator of metabolic health. Consumption of the branched-chain amino acid isoleucine is positively correlated with body mass index in humans, and reducing dietary levels of isoleucine rapidly improves the metabolic health of diet-induced obese male C57BL/6J mice. However, it is unknown how sex, strain, and dietary isoleucine intake may interact to impact the response to a Western Diet (WD). Here, we find that although the magnitude of the effect varies by sex and strain, reducing dietary levels of isoleucine protects C57BL/6J and DBA/2J mice of both sexes from the deleterious metabolic effects of a WD, while increasing dietary levels of isoleucine impairs aspects of metabolic health. Despite broadly positive responses across all sexes and strains to reduced isoleucine, the molecular response of each sex and strain is highly distinctive. Using a multi-omics approach, we identify a core sex- and strain-independent molecular response to dietary isoleucine, and identify mega-clusters of differentially expressed hepatic genes, metabolites, and lipids associated with each phenotype. Intriguingly, the metabolic effects of reduced isoleucine in mice are not associated with FGF21 – and we find that in humans plasma FGF21 levels are likewise not associated with dietary levels of isoleucine. Finally, we find that foods contain a range of isoleucine levels, and that consumption of dietary isoleucine is lower in humans with healthy eating habits. Our results demonstrate that the dietary level of isoleucine is critical in the metabolic and molecular response to a WD, and suggest that lowering dietary levels of isoleucine may be an innovative and translatable strategy to protect from the negative metabolic consequences of a WD.

## Introduction

Around the world, the prevalence of obesity has grown dramatically over the last few decades. In the United States, the age-adjusted percentage of adults who are either overweight or obese is now greater than 70% (NIDDK, 2021). Exposure to unhealthy “Western” diets begins early, and childhood obesity has steadily risen for about 50 years; about one-third of children in the United States are now overweight or obese (NIDDK, 2021). The health consequences of these effects are significant, as obesity is a risk factor not just for type 2 diabetes, but also for Alzheimer’s disease, cancer, cardiovascular disease, and hypertension (Dietary Guidelines for Americans, 2020-2025, 2020). While losing weight is a highly effective method to improve metabolic health, few individuals can consistently adhere to a reduced calorie diet; and while glucagon-like peptide-1 (GLP-1) agonists have recently been wildly successful at inducing weight loss through appetite suppression, they are expensive, have a wide-range of side effects, and their long-term consequences are unknown.

While flying in the face of conventional wisdom, over the past decade it has gradually become clear that calories derived from all macronutrient sources are not equivalent (Mihaylova et al., 2023). Accumulating evidence now supports a critical role for dietary protein in metabolic health. Dietary protein is often thought of as beneficial, promoting satiety, muscle growth, and blood sugar control, a view supported by some short-term human trials and studies of the elderly (Dong et al., 2013; Gannon et al., 2003; Kuzuya, 2022; Ribeiro et al., 2019; Seino et al., 1983). However, multiple prospective and retrospective cohort analyses suggest that consumption of dietary protein increases the risk of developing diabetes (Akter et al., 2021; Huang et al., 2020; Lagiou et al., 2007; Levine et al., 2014; Sluijs et al., 2010). Supporting this latter view of high dietary protein consumption as detrimental, two randomized clinical trials of protein restriction (PR) have found that PR reduces weight and adiposity and improves multiple aspects of glycemic control (Ferraz-Bannitz et al., 2022; Fontana et al., 2016). Research in mice, where PR has significant benefits for metabolic health (Maida et al., 2016; Solon-Biet et al., 2014; Solon-Biet et al., 2015), suggests that these seemingly paradoxical findings regarding the metabolic effects of dietary protein may be due to the ability of exercise to protect from the negative effects of dietary protein observed in sedentary rodents and the majority of humans (Trautman et al., 2023).

We hypothesized that the benefits of PR might result from decreased intake of specific dietary essential amino acids. Blood levels of the three branched-chain amino acids (BCAAs; leucine, isoleucine, and valine) have long been associated with diabetes, insulin resistance and obesity in humans and rodents (Batch et al., 2013; Connelly et al., 2017; Felig et al., 1969; Newgard et al., 2009), and we found that dietary restriction of the BCAAs is sufficient to promote glycemic control and reduce adiposity in both lean and diet-induced obese mice without restricting calorie intake (Cummings et al., 2018; Fontana *et al*., 2016; Richardson et al., 2021). BCAA-restricted mice had increased food intake that was more than offset by increased energy expenditure; similar beneficial results of restricting BCAAs have been observed in rats (White et al., 2016). Two short-term randomized clinical trials have observed metabolic benefits from BCAA restriction, including improved insulin resistance and white adipose tissue metabolism (Karusheva et al., 2019; Ramzan et al., 2020).

We recently determined that the metabolic benefits of BCAA restriction are driven specifically by reduced levels of isoleucine. Restriction of isoleucine is sufficient to promote metabolic health in lean mice, improving glucose tolerance through increased hepatic insulin sensitivity and decreasing adiposity by promoting the beiging of inguinal white adipose tissue (Yu et al., 2021). Further, the restriction of isoleucine is necessary for the full metabolic benefits of PR (Yu *et al*., 2021), and can robustly extend the healthspan and lifespan of adult male mice (Green et al., 2023). In diet-induced obese mice consuming a high-fat, high-sucrose Western diet (WD), the benefits of isoleucine restriction are even more profound; mice consuming a diet with reduced levels of isoleucine rapidly lose excess adipose mass, become glucose tolerant and insulin sensitive, and their fatty livers normalize (Yu *et al*., 2021). These results point to a key role of dietary isoleucine in the metabolic response to diet – and in particular, suggest that the dietary isoleucine in WD plays a key role in its negative effects on metabolic health.

We comprehensively examined the impact of dietary isoleucine level on the metabolic and molecular response to an unhealthy high-fat, high-sucrose WD. As sex and genetic background play an important role in the response to a variety of dietary interventions (Barrington et al., 2018; Browning et al., 2012; De Groef et al., 2021; Green et al., 2022; Mitchell et al., 2016; Roy et al., 2021), we challenged male and female mice from two different inbred strains, C57BL/6J and DBA/2J, with Western diets containing one of three different levels of isoleucine. Surprisingly, unlike the highly sex- and strain-specific metabolic responses of mice to PR (Green *et al*., 2022), we found that reducing dietary levels of isoleucine improved aspects of metabolic health such as body weight, energy expenditure, and glucose tolerance across all sexes and strains, albeit to different degrees, while aspects of metabolic health were impaired in all groups when dietary levels of isoleucine were elevated. Surprisingly, despite the similar effects of isoleucine on aspects of metabolic health, dietary isoleucine had diverse sex-specific and strain-specific effects on molecular pathways. Using a multi-omics approach, we identified a conserved sex and strain-independent molecular signature of isoleucine restriction in the liver of mice; we also identified gene-metabolite-lipid mega-clusters that correlate with key phenotypes, finding that the expression of the hormone FGF21 does not group with energy expenditure. While in humans BMI is positively associated with dietary isoleucine levels (Yu *et al*., 2021), we find that plasma FGF21 in humans is not significantly associated with dietary isoleucine. Finally, we find that animal-derived protein contains more isoleucine than plant-derived protein, and that there is diversity in BCAA levels between individual foods in different category types. Finally, we find that individuals surveyed in the National Health and Nutrition Examination Survey (NHANES) eating the healthiest diets consumed the lowest amount of isoleucine relative to their intake of dietary protein.

We conclude that isoleucine is a potent regulator of metabolic health across sexes and strains of mice, and that the dietary level of isoleucine is critical in determining the metabolic impact of a WD. Our results further suggest that consuming less isoleucine, which can be achieved in the context of a natural human diet through attentive food choices, may be an easy and translatable way to protect individuals from the seemingly ubiquitous exposure to Western diets.

## Results

### Dietary isoleucine levels are associated with WD-induced increases in body weight and fat mass

To examine the specific role of dietary isoleucine on the metabolic response to a WD, we designed a series of diets based on an amino acid-defined Western diet (WD Control) that we have previously utilized (Cummings *et al*., 2018; Yu *et al*., 2021), and which closely matches the macronutrient profile of the widely used natural source high-fat high-sucrose Western diet TD.88137. We designed diets with three different levels of isoleucine: the WD Control, a diet with a 67% reduction in isoleucine (WD Low Ile) relative to the WD Control, and a diet in which isoleucine is increased three-fold (WD High Ile) relative to the WD Control diet. These three diets are isocaloric with identical levels and sources of fat and carbohydrates, and the percentage of calories derived from AAs was kept constant by proportionally adjusting the amount of non-essential AAs. We also utilized a fourth diet group, which was fed an AA-defined diet with a composition similar to that of a natural chow diet (Ctrl AA); the amino acid profile of the WD Control AA and Ctrl AA diet are matched. The full composition of these diets can be found in **Table S1**.

We fed these diets to both male and female C57BL/6J (B6) and DBA/2J (DBA) mice for 3 months starting at 6 weeks of age (**Fig. 1A**). We saw an increase in food intake on WD Low Ile diets relative to the WD Control fed mice in all sex-strain groups, which was statistically significant in B6 and DBA males and trended higher in DBA females (p = 0.0638) and B6 females (**Fig. 1B**). Food consumption was not significantly altered in any group of mice by the WD High Ile diet (**Fig. 1B**).

**Figure 1.**
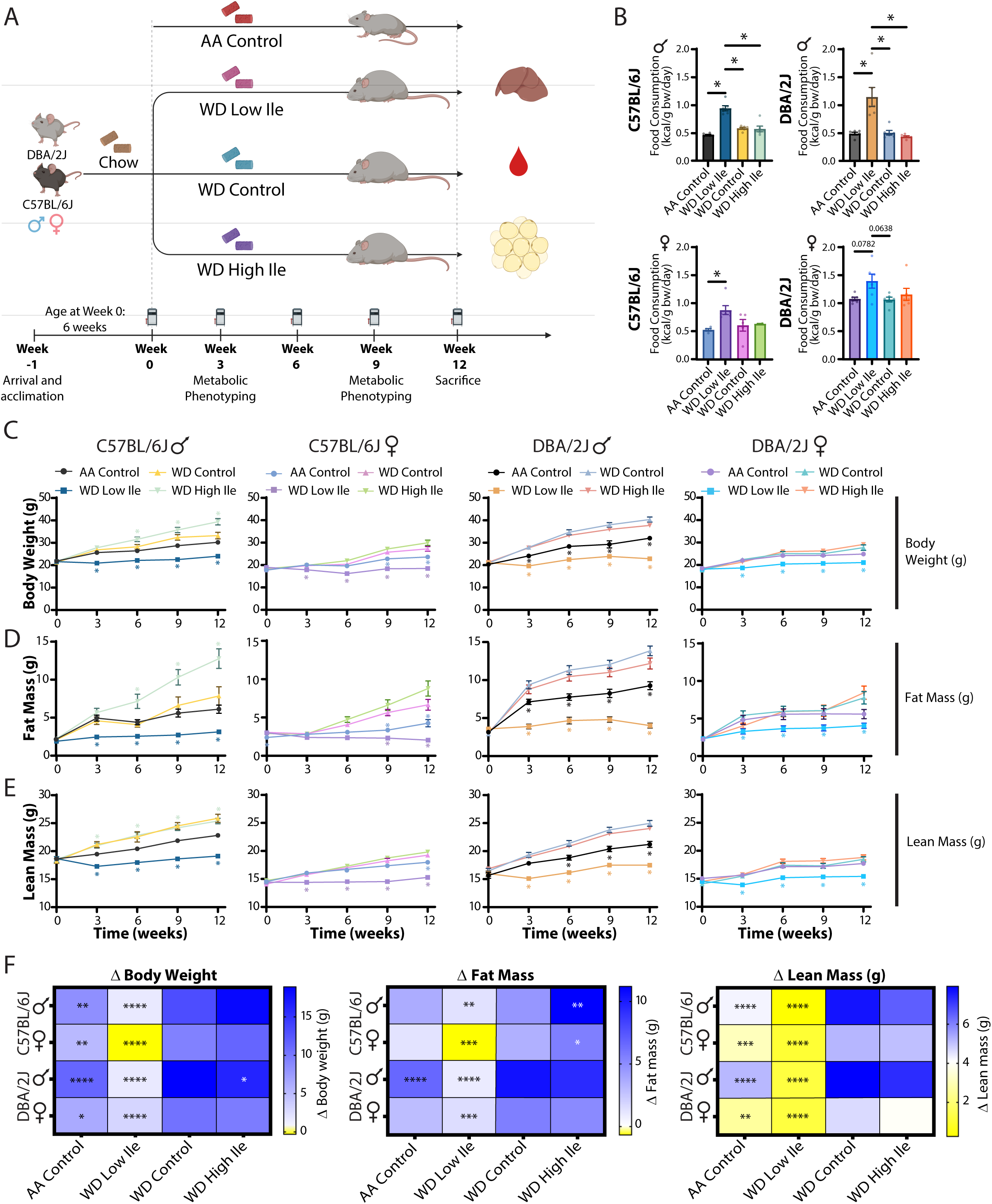
Reduced dietary isoleucine protects from WD-induced weight and fat gain. (A) Experimental design. (B) Food consumption normalized to body weight after 3 or 8 weeks on the indicated diets. (C-F) Body weight (C), fat mass (D) and lean mass (E) was tracking longitudinally, and change (Δ) in grams from the beginning to the end of the study was calculated (F). (B) n=2-12 mice per group. Statistics for the overall effects of diet represent the p value from a one-way ANOVA when compared to the WD Control diet; *p<0.05, Sidak’s post-test examining the effect of parameters identified as significant in the one-way ANOVA. (C-F) For longitudinal studies, statistics for the overall effects of time, diet, and the interaction represent the p value from a two-way RM ANOVA or residual maximum likelihood (REML) analysis conducted individually for each sex and strain. Each diet was compared to the WD Control diet; *p<0.05, Dunnett’s post-test examining the effect of parameters identified as significant in the two-way ANOVA. Data represented as mean ± SEM.

We tracked the weight and body composition of all groups for 12 weeks (**Figs. 1C-F**). Despite the increased food consumption of WD Low Ile-fed mice, all groups of mice consuming the WD Low Ile diet gained less body weight than WD Control-fed mice during the course of the experiment (**Figs. 1C, F**). In most groups, this was due to reduced accretion of both fat mass and lean mass by WD Low Ile-fed mice, although B6 females showed a net decrease of fat mass (**Figs. 1D-F**). The overall effect was that the WD Low Ile-fed mice of all groups had the smallest increase in adiposity during the course of the experiment, with B6 females showing an overall decline in adiposity after 12 weeks (**Supplemental Figs. 1A-D**). Conversely, B6 males fed a WD High Ile diet gained significant additional weight and accreted significant additional fat mass relative to WD Control-fed males; and an overall increase in adiposity was seen in WD High Ile-fed B6 males and females (**Fig. 1D and Supplemental Figs. 1A-B**). Altogether, mice eating diets with lower levels of Ile had reduced body weight, fat mass gain, and adiposity relative to mice eating diets with higher levels of isoleucine, while mice eating WD High Ile diets showed strain and sex-specific increases in weight and fat mass relative to WD Control-fed mice (**Fig. 1F**).

### Dietary isoleucine restriction protects from the negative effects of a WD on glycemic control

During the course of the experiment, we assessed the effect of isoleucine on glucose homeostasis of WD-fed mice. After three weeks on diet, we found that Ile restriction improved glucose tolerance in all mice, with WD Low Ile-fed mice having significantly reduced area under the curve (AUC) relative to WD Control-fed mice in all groups except DBA females (p=0.0543) (**Supplemental Fig. 2A**). After 9 weeks, all sexes and strains consuming the WD Low Ile diet had significantly improved glucose tolerance relative to WD Control-fed mice (**Fig. 2A**). Insulin sensitivity, as assessed by I.P. administration of insulin, was unaffected by dietary Ile in both male and female B6 mice; in male and female DBA mice, lower levels of Ile were associated with improved insulin sensitivity (**Fig. 2B and Supplemental Fig. 2B**).

**Figure 2.**
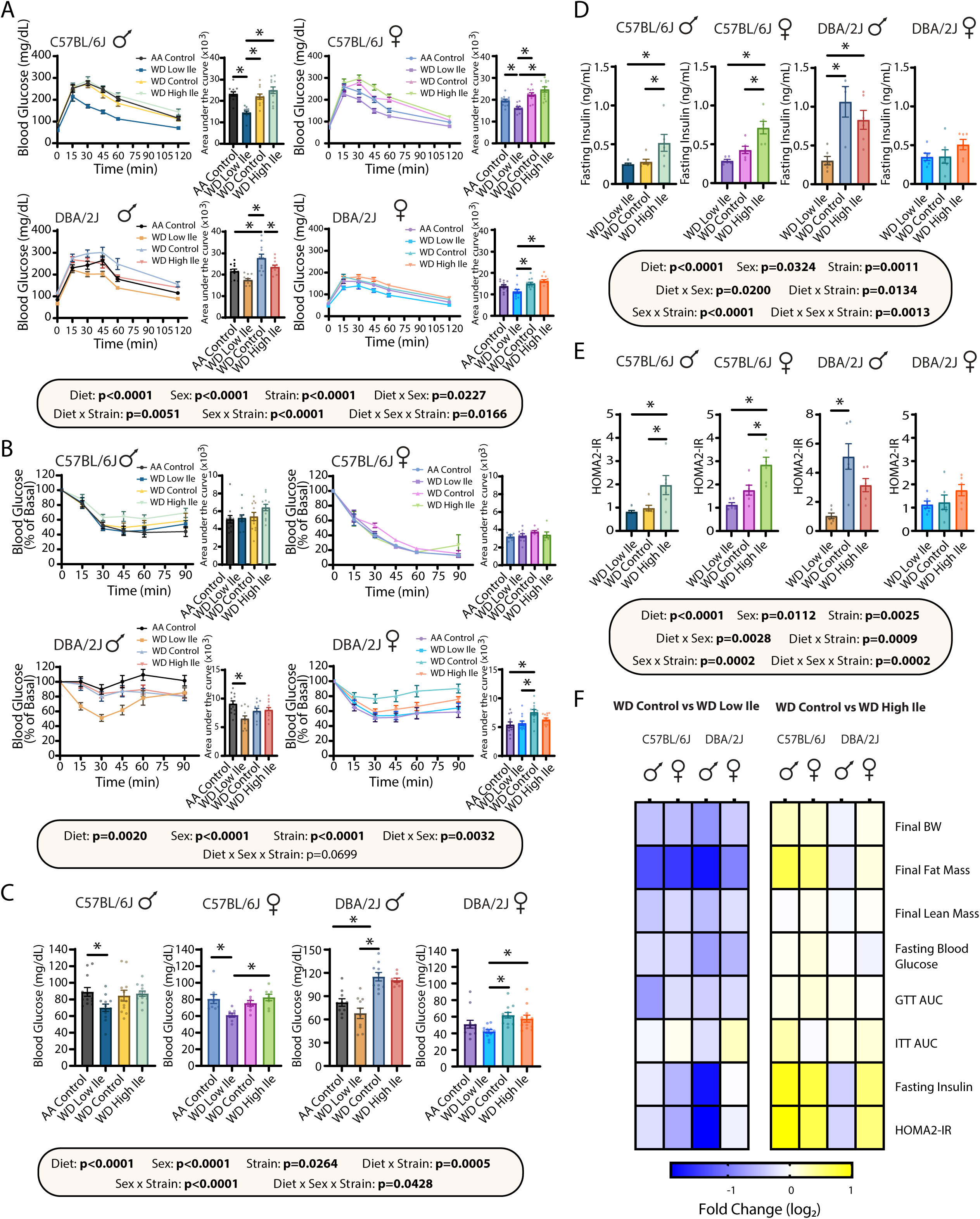
Dietary isoleucine is negatively associated with glycemic control in WD-fed mice. (A-B) Glucose (A) and insulin (B) tolerance tests in mice fed the indicated diets for 9 weeks and 10 weeks, respectively, with quantified area under the curve (AUC). (C-E) Fasting blood glucose (C) and insulin (D) levels were determined and HOMA2-IR (E) was calculated after 12 weeks on the indicated diets. (F) Heatmap of the effect of isoleucine on body composition and glucose homeostasis; log_2_ of the fold change throughout the experiment is plotted. (B-F) n=5-12 mice per group. Statistics for the overall effects of diet, sex, and strain represent the p value from a three-way ANOVA; *p<0.05, Tukey’s test post ANOVA for each sex/strain group shown. Data represented as mean ± SEM.

We also collected blood to measure fasting blood glucose and insulin levels and performed a homeostasis model assessment for insulin resistance (HOMA2-IR) (Geloneze et al., 2009; Mather, 2009). In almost all groups, fasted blood glucose following an overnight fast was significantly lower in WD Low Ile-fed mice relative to mice fed either WD Control or WD High Ile diets (**Fig. 2C**), with smaller differences between groups observed following a 4-hour daytime fast (**Supplemental Fig. 2C**). Across all sexes and strains, fasting insulin levels were correlated with dietary Ile levels, with the highest levels generally observed in WD High Ile fed mice and the lowest in WD Low Ile-fed mice, though these differences did not reach statistical significance in DBA females (**Fig. 2D**). In B6 mice, increased dietary levels of Ile were associated with higher insulin resistance as assessed by HOMA2-IR, while there was a trend toward lower HOMA2-IR in WD Low Ile-fed B6 females and a significantly lower HOMA2-IR in DBA males (**Figs. 2E-F**). We also calculated HOMA2-%B, which indicates pancreatic beta cell function; we observed an overall decrease in beta cell function in WD High Ile-fed males, which was statistically significant in B6 males (**Supplemental Fig. 2D**). Finally, we visualized the impact of dietary Ile levels on body composition and glucose homeostasis using heatmaps. As shown in **Fig. 2F**, reducing dietary isoleucine generally promoted metabolic health, improving parameters related to both body composition and glycemic control across sexes and strains, while increasing dietary isoleucine levels conversely impaired metabolic health, especially in B6 males and females.

### A low isoleucine diet increases energy expenditure, with strain and sex-dependent effects on the FGF21-UCP1 axis

We hypothesized that the effects of dietary isoleucine on weight and body composition were likely mediated by changes in energy balance. We utilized metabolic chambers to evaluate energy balance, analyzing energy expenditure and spontaneous activity. Based on our previous experiments with isoleucine restriction in diet-induced obese B6 males, we expected that the WD Low Ile diet would increase energy expenditure; in agreement with our prediction, we observed a robust increase in energy expenditure in B6 males and females during both the light and dark cycles (**Fig. 3A** and **Supplemental Fig. 3A**). Energy expenditure was also increased in the WD Low Ile-fed DBA mice, reaching statistical significance as compared to WD Control-fed mice during the dark cycle in DBA females; in DBA males, WD Low Ile-fed males had higher energy expenditure than WD High Ile-fed males during the dark cycle (**Fig. 3A** and **Supplemental Fig. 3A**). In both the light and dark phases, there was a strong effect of diet on energy expenditure. In the light phase, there were significant interactions between diet and strain as well as sex and strain on energy expenditure; in the dark phase, there was a significant effect of sex (**Fig. 3A**). Ile restriction has previously been shown to increase the respiratory exchange ratio (RER) of B6 males on a low-fat diet, suggesting an increased utilization of carbohydrates. Surprisingly, here we observed that RER decreases in WD Low Ile-fed B6 males relative to WD Control-fed mice, suggesting an increase in fat oxidation, which reached statistical significance during the light cycle; we also observed a similar effect of Ile restriction in DBA males, reaching statistical significance during the dark cycle (**Supplemental Fig. 3B**). In contrast, the RER of B6 and DBA females decreased as dietary Ile increased, which was statistically significant in B6 females in the dark cycle (**Supplemental Fig. 3B**). Overall, in the light phase we observed significant effects of diet, sex, and strain, as well as a significant interaction between diet and sex on RER, and found that there was a significant effect of diet on RER in the dark phase (**Supplemental Fig. 3B**).

**Figure 3.**
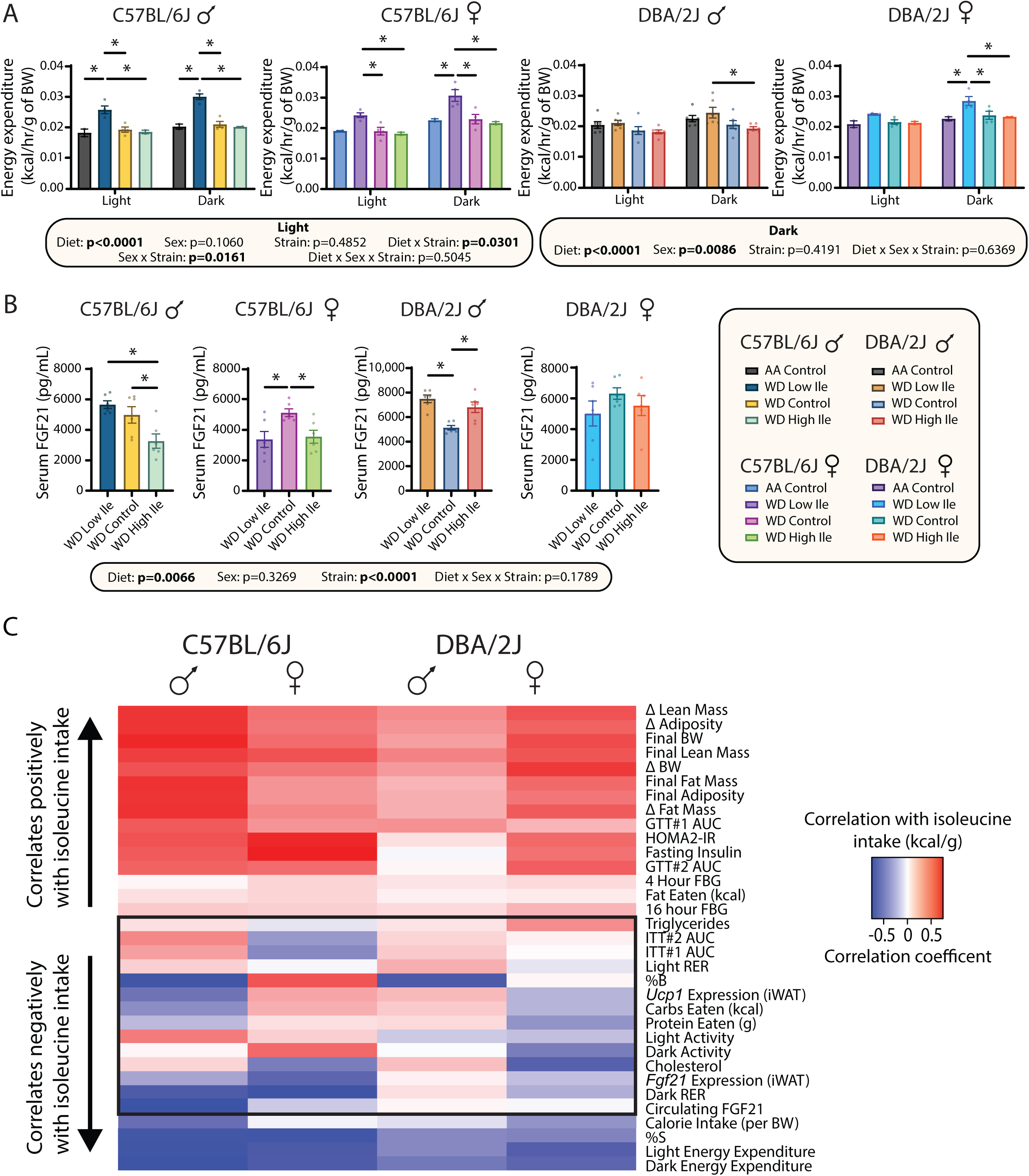
Dietary isoleucine is negatively associated with energy expenditure, but not FGF21, in WD-fed mice. (A) Energy expenditure per gram of body weight. (B) Circulating FGF21. (C) Phenotypic and molecular measurements correlated with consumption of isoleucine (kcals) in each mouse (Pearson’s correlation) and clustered (hierarchical clustering). Phenotypic measurements that do not cluster as well appear in the middle of the correlation plot (black outline). n=3-7 mice per group. Statistics for the overall effects of diet, sex, and strain represent the p value from a three-way ANOVA; *p<0.05, Tukey post-test examining the effect of parameters identified as significant in the one-way ANOVA. Data represented as mean ± SEM.

The increased energy expenditure of B6 males on a PR diet is mediated by fibroblast growth factor 21 (FGF21), a key energy balance hormone that is induced by PR in both humans and rodents, and which promotes the beiging of white adipose tissue (Flippo et al., 2020; Hill et al., 2019; Laeger et al., 2014). Ile restriction similarly induces FGF21 and induces the beiging of white adipose tissue in B6 males consuming a low-fat diet, increasing energy expenditure via a partially *Fgf21*-dependent pathway (Yu *et al*., 2021). However, we recently found that the effect of PR on FGF21 is dependent upon sex and strain, and that FGF21 may not be required for the metabolic response to PR in females (Green *et al*., 2022).

To better understand the role of FGF21 in the response to dietary isoleucine, we examined the effect of dietary Ile level on the FGF21-UCP1 axis across sexes and strains. In B6 males, we observed that FGF21 levels are significantly lower in WD High Ile-fed B6 males than in B6 males fed either the WD Control or WD Low Ile diets (**Fig. 3B**). However, this association with Ile level was not observed in other sexes and strains; in B6 and DBA females, the highest level was found in WD Control-fed animals, while in DBA males, FGF21 was elevated in both WD Low Ile-fed and WD High Ile-fed mice relative to WD Control-fed mice (**Fig. 3B**).

Next, we examined relative *Fgf21* and thermogenic and lipolytic gene expression in inguinal white adipose tissue (iWAT). We were surprised to see that unlike active FGF21 in circulation, relative *Fgf21* expression in iWAT was largely unchanged in all groups, except for DBA females (**Supplemental Fig. 3C**). Both high and low Ile-fed DBA females displayed significantly reduced *Fgf21* compared to the WD Control (**Supplemental Fig. 3C**).

In the context of a low-fat diet, we have previously observed that isoleucine restriction increases the expression of Uncoupling protein 1 (*Ucp1)* and other thermogenic and lipolytic genes, including cell death-inducing DNA fragmentation factor alpha-like effector A (*Cidea*), elongation of very long chain fatty acids protein 3 (*Elovl3*), Peroxisome proliferator-activated receptor gamma (*Pparg*), hormone-sensitive lipase (*Lipe*) and fatty acid synthase (*Fasn*) (Yu *et al*., 2021) (**Supplemental Fig. 3C, Table S10**). Here we show only B6 males displayed a robust increase in *Ucp1, Pparg, Fasn, Lipe and Cidea*, showing clear sex and strain-dependent FGF21-UCP1 signaling when consuming a WD. (**Supplemental Fig. 3C, Table S10**). Intriguingly, DBA males under the WD Control exhibited increased signaling of these genes compared to both high and low Ile diets (**Supplemental Fig. 3C, Table S10**). These data suggest that isoleucine levels regulate energy expenditure largely independently of the FGF21-UCP1 axis and iWAT beiging.

In addition to energy expenditure parameters, we examined how dietary isoleucine levels impacted cholesterol and triglycerides. Despite evidence that in response to a Western style diet, B6 mice display a greater rise in circulating cholesterol and triglycerides than DBA animals (Zhu et al., 2009), we observed roughly the same levels of plasma cholesterol in all groups, regardless of diet (**Supplemental Fig. 4A**). WD Low Ile-fed mice had significantly lower plasma triglycerides in DBA males, and we observed a similar effect in B6 females (p=0.0677) (**Supplemental Fig. 4B**).

### Metabolic phenotypes and hepatic -omics highlight sex and strain-dependent differences in the response to dietary Ile

We next used multivariate analysis to comprehensively identify sex- and strain-dependent responses to dietary Ile. We correlated the Ile intake of each individual mouse on the WD Low Ile, WD Control, and WD High Ile diets with 33 phenotypic and molecular measurements obtained from each animal for each strain and sex, and used hierarchical clustering to determine patterns of change relative to Ile intake (**Fig. 3C**). Overall, there was high similarity in the response to altered isoleucine intake across sexes and strains, with Ile intake positively correlating with body weight, fat mass, adiposity, and AUC during a glucose tolerance test, and Ile intake correlating negatively with calorie intake and energy expenditure. In the middle of the plot (black outline, **Fig. 3A**) are phenotypes which tend to vary the most in response to sex and strain; these include insulin sensitivity, activity, and RER as well as calories of carbohydrate consumed.

We performed a principal component analysis (PCA) with the phenotypic data, plotting males and females separately (**Supplemental Figs. 4C-D**). We observed a significant overlap between WD Low Ile-fed B6 and DBA males, indicating a strong effect of diet over strain in Ile restriction. In contrast, B6 WD Control-fed males largely overlapped with WD High Ile-fed males, and DBA WD Control-fed males overlapped with WD High Ile-fed males, suggesting that strain was more responsible for the variation between groups than diet when Ile content is normal or high (**Supplemental Fig. 4C**). This pattern was not as strong in females, with only partial overlap of B6 and DBA females fed the WD Low Ile diet, and the groups appearing to cluster primarily by strain (**Supplemental Fig. 4D**). The phenotypes with the greatest contribution to the spread of the PCA in males were fasting blood glucose, final fat mass/adiposity, and circulating FGF21 (**Supplemental Fig. 4E**), while in females the main drivers included energy expenditure, HOMA2-IR, RER, and food consumption (**Supplemental Fig. 4F**).

Due to the central role of the liver in maintaining metabolic homeostasis, we performed a detailed molecular analysis of the livers of WD Low Ile, WD Control, and WD High Ile-fed mice of both sexes and strains, performing transcriptional profiling as well as metabolomic and lipidomic analysis. We first analyzed the effects of dietary Ile on each type of data separately.

At the transcriptional level, we saw many sex- and strain-dependent effects on gene transcription (**Table S5**) and we identified significantly altered pathways using KEGG pathway enrichment analysis. As shown in **Figure 4A**, there are clear effects of sex and strain on the molecular response to dietary Ile. Comparing the WD Low Ile diet to the WD Control diet, we found two significantly downregulated and 16 significantly upregulated pathways in B6 males, many of metabolic interest, such as “Glutathione metabolism,” “Lysosome,” and “Terpenoid backbone biosynthesis” (**Fig. 4A**). However, many more and different pathways were altered in B6 females and DBA males, while very few pathways were significantly altered in DBA females (**Fig. 4A**). Comparing the WD High Ile diet and the WD Control diet, robust changes were observed only in B6 males, while neither B6 nor DBA females had significant changes in KEGG pathways in response to a WD High Ile diet (**Supplemental Fig. 5A**).

**Figure 4.**
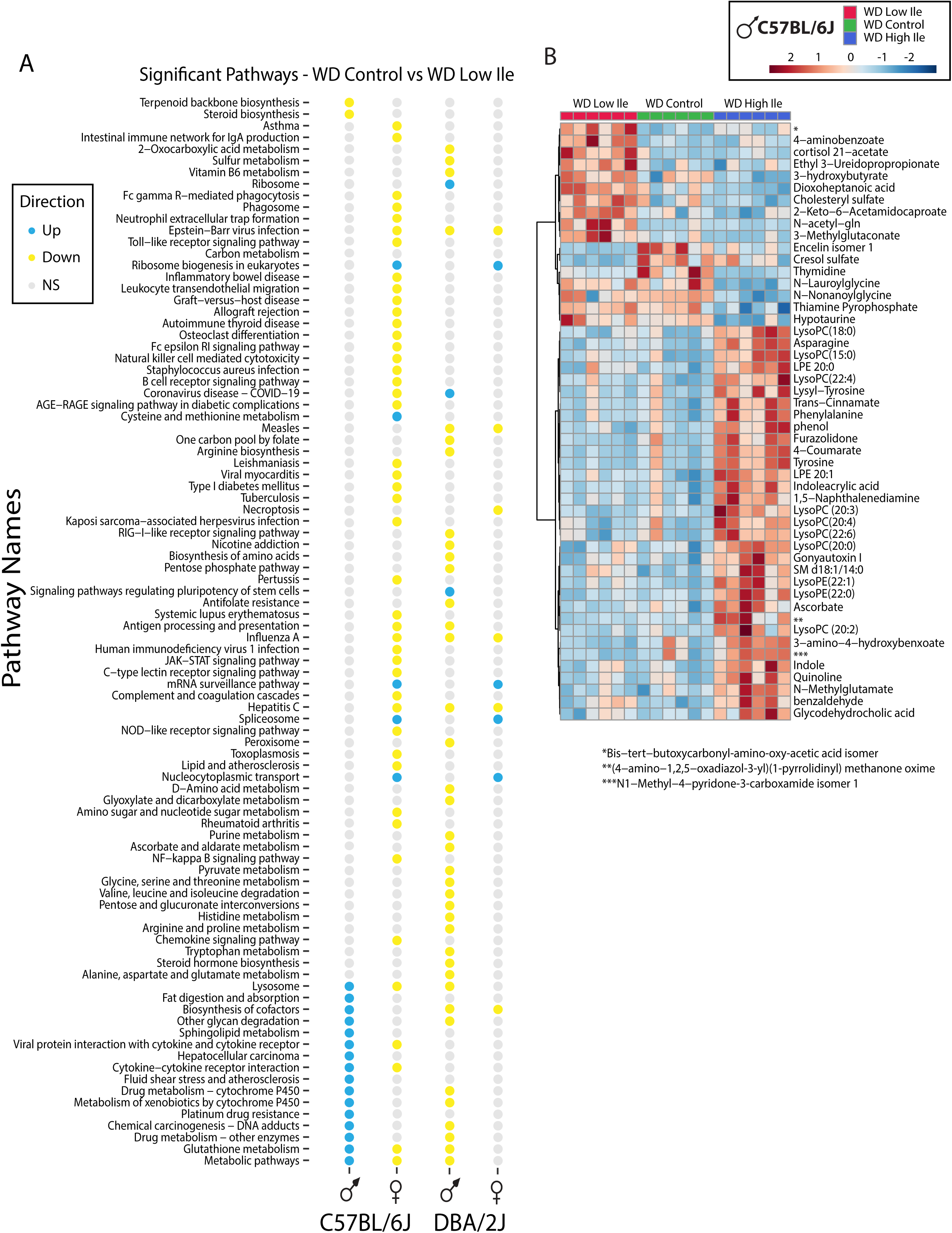
Altering the isoleucine content of a WD has sex and strain-specific effects on the hepatic transcriptome and metabolome. (A) KEGG enrichment pathway analysis of significantly altered genes in WD Low Ile-fed mice relative to WD Control-fed mice in each sex and strain (adjusted p<0.05). (B) Top 50 hepatic metabolites altered in response to dietary isoleucine in C57BL/6J male mice consuming a Western diet.

Both of these outcomes were in agreement with what we observed during our metabolic phenotyping; that is, strong changes in males, especially of the B6 background, and muted responses in females, especially DBAs. Intriguingly, insulin resistance and the insulin signaling pathway were the only pathways that were significantly increased in DBA males in response to a WD High Ile diet (**Supplemental Fig. 5A**), which is consistent with our ITT results, but not HOMA-IR findings (**Figs. 2B, 2E**). There were no overlapping genes between WD High Ile-fed B6 and DBA male mice (**Supplemental Fig. 5C**).

Next, we analyzed the hepatic metabolome. Though there were clear patterns present between the diet groups within the same sex and strain, the significant metabolites altered in different sexes and strains were quite different (**Fig. 4B** and **Table S6)**. In B6 animals, the impact of the WD High Ile diet on metabolites was limited to males, as additional dietary Ile did not make much of an impact on liver metabolites in females (**Fig. 4B and Supplemental Fig. 6A**). In DBA mice, the WD High Ile diet had the strongest effect on hepatic metabolites changes in males, while in females, different metabolites were increased by isoleucine restriction or supplementation (**Supplemental Figs. 6A-C**).

We also identified sex- and strain-dependent changes at the lipid level, with significant changes in pathways altered by both reduction and supplementation of dietary isoleucine (**Supplemental Fig. 6D**). Consistent with our transcriptomics and metabolomics results, there was little conservation of lipid species and pathways that were significantly altered between sex and strains; lipidomic analysis revealed that no single pathway was simultaneously altered in all four groups in response to either diet (**Supplemental Fig. 6D**). Some trends did emerge, such as the shared downregulation of polyunsaturated fatty acids, glycerophosphoglycerols, and fatty acids with more than 3 double bonds in B6 males and females in the WD Control vs WD Low Ile analysis. There were also instances of two groups sharing a significantly altered pathway in the WD High Ile analysis, but overall there were far fewer lipids and lipid pathways changed in response to elevated Ile (**Supplemental Fig. 6D**).

### Multiomic identification of a conserved molecular signature of isoleucine restriction

In order to identify a conserved molecular response to dietary isoleucine, we generated Venn diagrams of significantly up and down-regulated genes, lipids, and metabolites in response to WD Low Ile (**Fig. 5A-F, Table S8**). and the genes of WD High Ile diets (**Supplemental Fig. 5B-C**). In response to a WD Low Ile diet, we observed increased expression in all sexes and strains of genes related to the circadian CLOCK-BMAL transcription complex (**Fig. 5A**), and decreased expression in all sexes are strains of genes involved in nitrogen metabolism, metabolic pathways, and protein processing in the endoplasmic reticulum (ER) (**Fig. 5B**).

**Figure 5.**
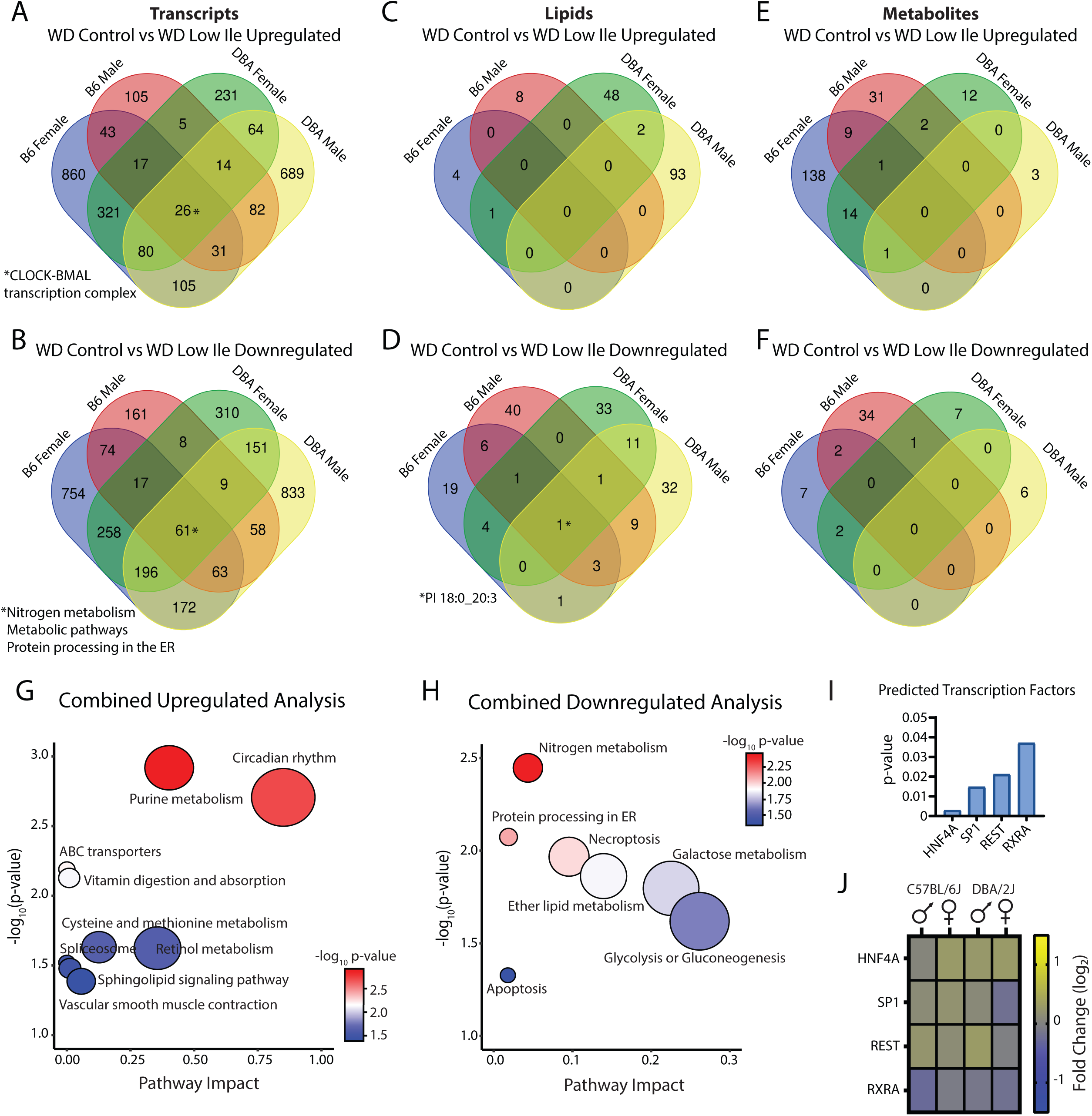
Analysis across sexes and strains identifies a conserved molecular response of the liver to restriction of dietary isoleucine. (A-F) Venn diagrams of the genes (A-B), lipids (C-D), and metabolites (E-F) significantly altered by a WD Low Ile diet in the livers of mice of the indicated strain and sex. (G-H) Metaboanalyst was used to identify pathways altered in the shared upregulated (G) and downregulated (H) genes and lipids identified in A-F. (I-J) Predicted transcription factors driving the genes altered in all sexes and strains (I) and their corresponding expression level (J).

There were no conserved upregulated lipids, but there was one lipid downregulated in all groups, a phosphatidylinositol (PI 18:0_20:3) (**Fig. 5C-D**). Lastly, there were no conserved upregulated metabolites, but B6 females and DBA males and females all had significantly lower levels of glycerylphosphorylethanolamine (**Fig. 5E-F**). To further evaluate a conserved molecular hepatic signature by Ile restriction, we combined the genes, metabolites and lipids conserved in **Fig. 5A-F** and used Metaboanalyst to identify significantly up- and downregulated pathways (**Fig. 5G-H, Table S7**). We find that in response to a WD Low Ile diet, circadian rhythm and purine metabolism pathways are strongly upregulated in the liver, as well as ABC transporters and vitamin digestion and absorption (**Fig. 5G**). Conversely, nitrogen metabolism, protein processing in the ER, necroptosis, ether lipid metabolism, galactose metabolism, glycolysis or gluconeogenesis, and apoptosis pathways are all downregulated (**Fig. 5H**).

Using the conserved significantly up and downregulated genes from our transcriptomics data, we utilized the MAGIC transcription factor prediction database (Roopra, 2020), where we identified 4 significantly enriched transcription factors (**Fig. 5I**). Based on our transcriptomics data, the majority of these genes change expression less than 2 fold – suggesting that any changes driven by these TFs are mediated by post-translational modifications, nuclear localization, and/or changes in DNA binding (**Fig. 5J**).

### Multi-omics analysis reveals that dietary Ile is a potent driver of molecular change

To examine the relationship between molecular changes and whole-organism physiology and metabolism induced by changes in dietary Ile, we constructed a Spearman’s Rank Order Correlation matrix using phenotypic data as well as hepatic transcriptomic, lipidomic and metabolomic data, a technique we have previously utilized to gain insight into the effects of dietary macronutrients (Green *et al*., 2022; Green *et al*., 2023). Statistically significant changes in gene expression, lipids and metabolites for each group of mice were concatenated with phenotypic data, and Spearman’s correlation was used to calculate coefficients that were then rendered by hierarchical clustering (based on 1-correlation coefficient between all molecules) to produce 9 distinct clusters (**Table S9**).

We used KEGG enrichment for the genes and metabolites in each cluster, and LiON to identify lipid ontology enriched terms in each cluster, and related these to the corresponding cluster phenotypes. While these results are based on correlations and are therefore not necessarily causative, they are consistent with a critical role for Ile, and link specific molecular and metabolic processes in the liver directly to the robust phenotypic response to changes in the level of dietary Ile. In cluster 1, we saw that energy expenditure relative to body weight correlated with “Isoleucine biosynthesis”, but not circulating FGF21, which presented in cluster 8 (**Fig. 6A, clusters 1, 8**). This again suggests that the increased energy expenditure of WD Low Ile-fed mice may be due to a non-canonical and FGF21-independent pathway. Indeed, FGF21 did not cluster with other phenotypes or KEGG terms related to metabolism beyond “metabolic process.” (**Fig. 6C, cluster 8**).

**Figure 6.**
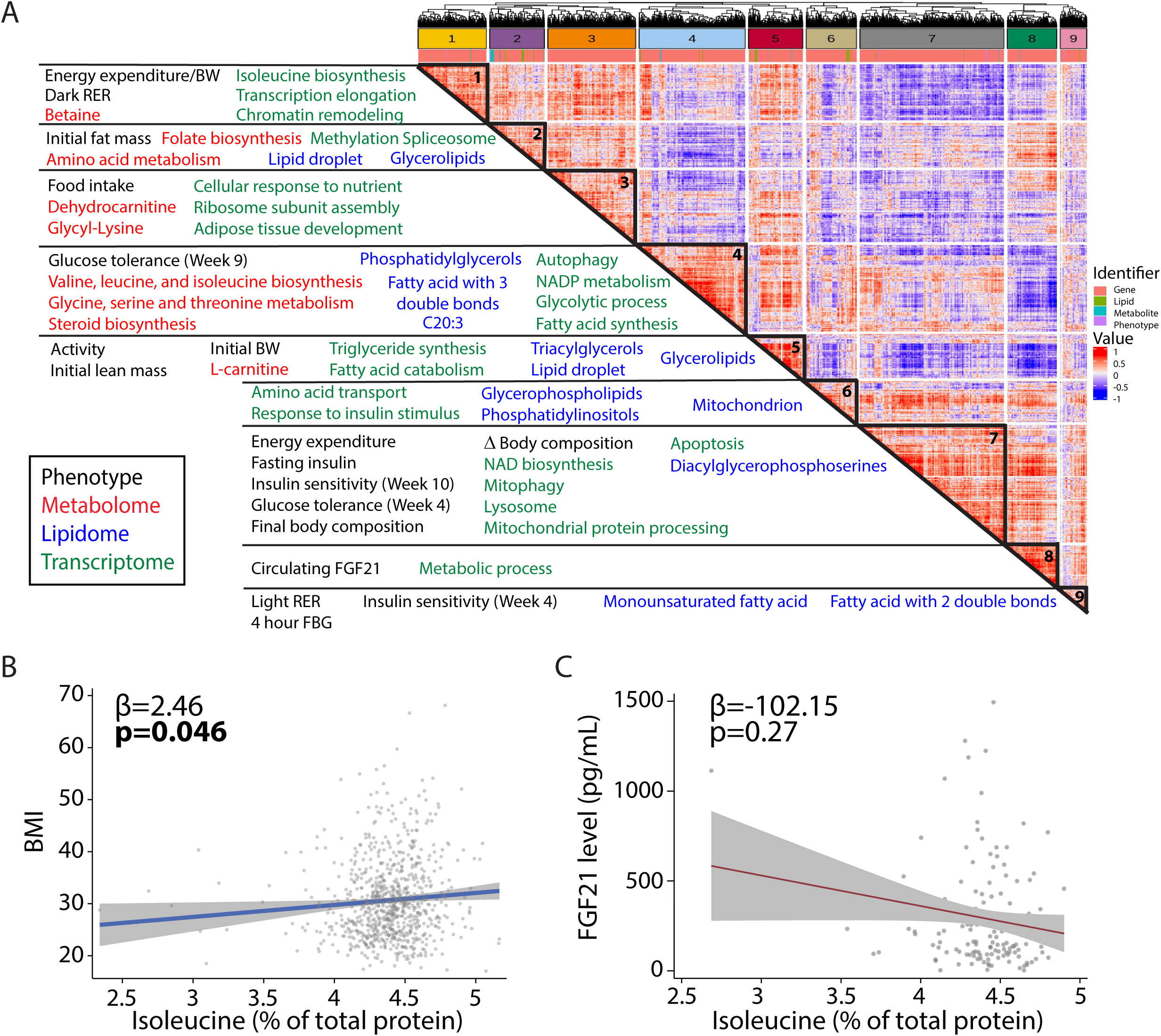
Integrated multiomic analysis links key phenotypic responses to reduced isoleucine with molecular and metabolic processes. (A) Spearman’s Rank Order Correlation matrix utilizing hepatic metabolomics, lipidomics, and transcriptomics data to correlate with phenotypes. n=6 per group. Isoleucine restriction produces changes across phenotypes that correlate with genes, metabolites, and lipids. Spearman’s rank order correlation matrix of significant observations, including phenotypic (black), transcriptomic (green), metabolomics (red), and lipidomic (blue) changes between WD Control and WD Low Ile diet groups. Hierarchical clustering identified 9 mega-clusters (outlined in black; **Table S9**). Enriched pathways listed in **Table S5**; phenotypes and pathways of interest in each cluster are highlighted. (B) Association between BMI and percent of total protein from Ile from the SHOW study (n = 788, shaded area represents 95% CI); previously published in (Yu *et al*., 2021). (C) Association between circulating FGF21 and dietary isoleucine intake (n = 140, shaded area represents 95% CI).

Not surprisingly based on our glucose homeostasis *in vivo* data, BCAA metabolism clustered with glucose tolerance, as well as autophagy, glycolysis, and fatty acid synthesis (**Fig. 6A, cluster 4**). Interestingly, the KEGG term “Lysosome” is associated with insulin sensitivity in cluster 7, suggesting that isoleucine may act to regulate blood sugar in part via lysosomal signaling. While not a conventional pathway, this possibility is consistent with emerging evidence for a role of the lysosome as a key regulator of glucose metabolism (Mancini et al., 2023) (**Fig. 6A, cluster 7**). Mitophagy, NAD+ and mitochondrial protein processing are also present in this cluster (**Fig. 6C, cluster 7**).

### Dietary isoleucine levels in humans are associated with BMI, but not FGF21 levels

As we have previously described, when we examined associations between estimated isoleucine intake relative to total protein and BMI (kg/m^2^) among a randomly selected, cross-sectional population-based sample of 788 adults, we found that after adjustment for confounding factors, an increase in the intake of dietary isoleucine relative to total protein of a single percentage point—for example, from 4% to 5% of protein—is associated with a 2.46 unit increase in BMI (p = 0.046) (**Fig. 6B**, (Yu *et al*., 2021)). To determine if FGF21 levels in humans are correlated with isoleucine, we examined stored plasma from 140 of these same subjects. Here we see that consistent with our mouse findings across sexes and strain, human plasma levels of FGF21 when adjusted for confounding factors are not associated with dietary levels of isoleucine relative to total protein (p=0.27) (**Fig. 6C**).

### Healthy human diet patterns are lower in relative Ile

We next investigated the contribution of dietary isoleucine to human nutrition and health using food item databases and cross-sectional dietary reporting data from the National Health and Nutrition Examination Survey (NHANES). We first looked at relative proportions of isoleucine across food item groups using the United States Department of Agriculture (USDA) National Nutrient Database for Standard Reference, Release 28. When we compared the amino acid composition of plant vs. animal-based food items, we observed that animal-based items have higher proportional isoleucine content (**Fig. 7A**). While the isoleucine content of animal-based foods followed a largely unimodal distribution, plant-based items had a proportionally lower isoleucine content but showed greater variability, following a bimodal distribution with a small high proportion sub-group (**Fig. 7A**). Further sub-division of food items revealed that several animal-based food groups had the highest proportional abundance of isoleucine, including beef, fish, pork and poultry (**Fig. 7B**). Interestingly, fruits had the lowest proportional isoleucine abundance followed by nuts and seeds and cereal grains, while legumes tended to have the highest relative isoleucine abundance among plant-based items (**Fig. 7B**). The same patterns in food items were observed when we examined the other two BCAAs, leucine and valine, except that isoleucine tended to have lower relative abundance in cereal grains (**Supplemental Fig. 7A**).

**Figure 7.**
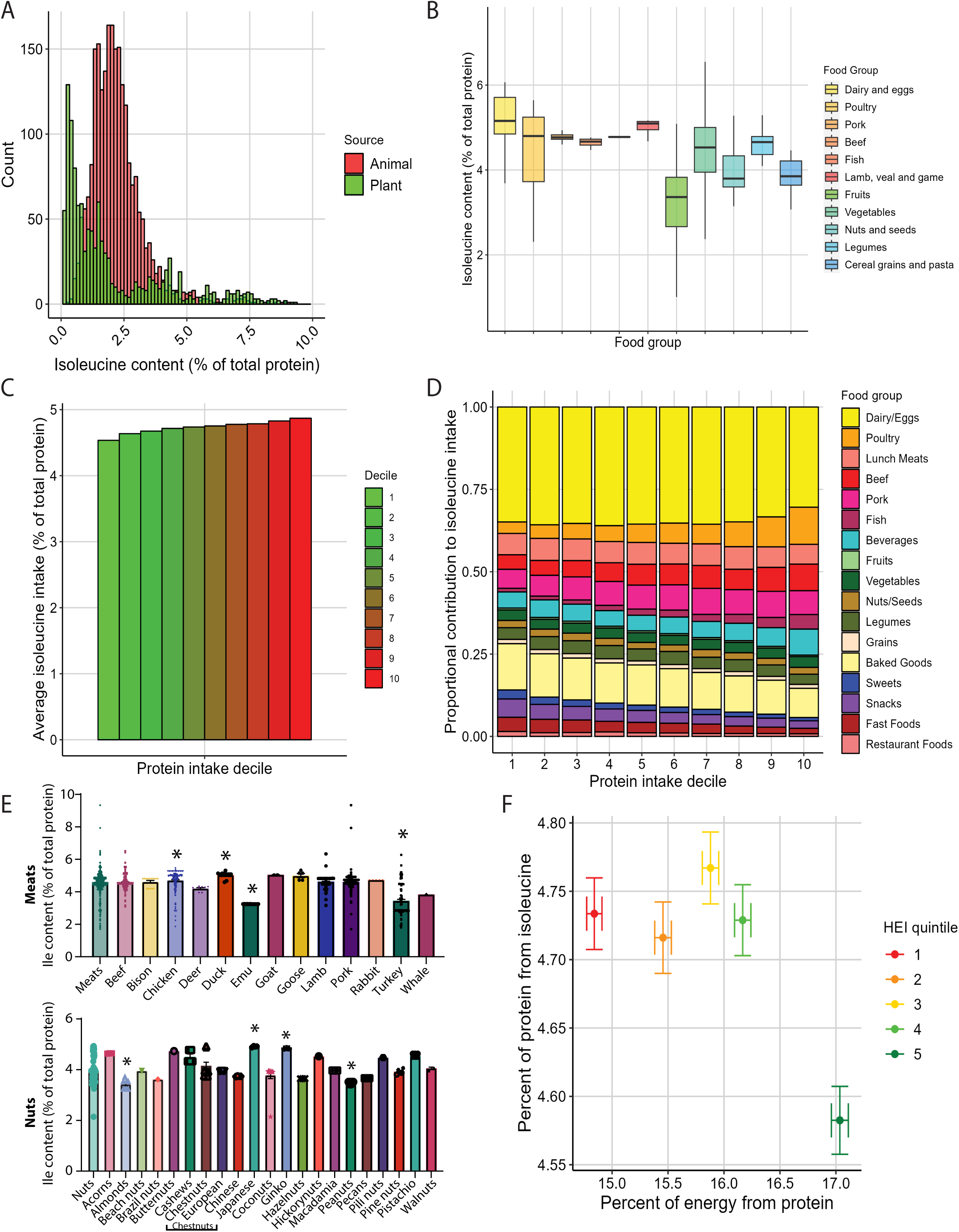
People eating the healthiest diets consume relatively less isoleucine. (A-C) Relative isoleucine content in animal versus plant foods (A), by food group (B), and per decile of protein intake (C). (D) Proportional contribution of isoleucine intake per decile of protein intake by food group. (E) Relative isoleucine content in commonly consumed meats and nuts. *p<0.05, Dunnett’s post-test vs. Meats or Nuts. (F) Relative isoleucine per protein intake and HEI quintile scores. Data represented as mean ± SEM.

We next looked at isoleucine consumption across dietary patterns using NHANES data. When stratifying individuals by decile of protein energy intake (% of kcals), a clear trend emerged with lower protein consumers eating a lower relative proportion of isoleucine (**Fig. 7C**). Interestingly, this trend was not evident with respect to leucine and valine, and we considered the possibility that the effect of isoleucine was driven by the lower relative abundance of isoleucine in grain-based food items (**Supplemental Fig. 7B**). Identifying how dietary food groups varied with protein intake, we find that lower protein consumers obtained a larger relative amount of isoleucine from items including baked goods and grain, while the highest protein consumers obtained a larger relative amount of their daily isoleucine from poultry, beef and beverages (including powdered protein drinks) (**Fig. 7D**).

Based on this initial data, we hypothesized that there would also be significant variation in the isoleucine content of individual foods in each category. As substitutions within a category – e.g., one vegetable for another – may be easier to incorporate into standard diets, we analyzed individual foods for the relative percentage of their isoleucine content (**Fig. 7E, Supplemental Fig. 7C**). Within the category of Meats, we found that emu and turkey had some of the lowest isoleucine content (**Fig. 7E**), while within Fruits, grapes and grapefruits were significantly lower than the other fruits (**Supplemental Fig. 7C**), and within Nuts, almonds and peanuts contained the lowest relative amount of isoleucine (**Fig. 7E**). Although there were no specific vegetables that were significantly lower in isoleucine, several individual vegetables, including carrots, chard, and lima beans had very high levels of isoleucine (**Supplemental Fig. 7C**).

Finally, we stratified participants based on indices of dietary health. Interestingly, when participants were stratified by healthy eating index (HEI) score, we found that despite participants in the highest HEI quintile having the greatest protein intake, they also had the lowest proportional intake of isoleucine, which may be due to relatively higher intake of foods with lower isoleucine levels (**Fig. 7F**).

## Discussion

Obesity is a growing problem throughout the world; not only are many adults overweight or obese, so are an increasing percentage of children and teenagers. In the United States, 19.7% of those 2-19 years old are obese, with many additional people in this age range likely overweight (National Health and Nutrition Examination Survey 2017–March 2020 Prepandemic Data Files Development of Files and Prevalence Estimates for Selected Health Outcomes, 2021). While a number of causes contribute to the early onset of obesity, lifelong consumption of an unhealthy Western diet is likely a critical factor. Intervening in the context of already established diet-induced obesity is clearly important; perhaps even more critical, however, is understanding how to prevent the development of obesity in the first place.

Based on emerging evidence of the regulation of metabolic health by the dietary branched-chain amino acid (BCAA) isoleucine (Green *et al*., 2023; Mihaylova *et al*., 2023; Solon-Biet *et al*., 2014; Yu *et al*., 2021), and a growing awareness that sex and genetic background impact the response to diet (Barrington *et al*., 2018; Green *et al*., 2022), we examined here the impact of sex and strain on the response to dietary intake of isoleucine. We find that lower levels of isoleucine consumption in the context of a Western diet promote metabolic health in all groups, including both sexes of C57BL/6J and DBA/2J mice. However, the size of this effect varied across groups; the greatest metabolic benefits of reduced isoleucine intake were observed in C57BL/6J males, with some of the smallest effects in DBA/2J females. We show that restriction of isoleucine has sex- and strain-dependent and independent effects on hepatic metabolism, and identify a conserved molecular signature of isoleucine restriction. Finally, we show that blood levels of FGF21 are not associated with dietary isoleucine levels in mice or humans, and examine how dietary intake of isoleucine varies with food source and dietary patterns.

Our studies here were conducted in mice consuming a high-fat, high-sucrose WD as they transition from adolescents (6 weeks old) to young adults (18 weeks old) – precisely aligning with the period in humans where obesity often begins (Flurkey et al., 2007). Surprisingly, our findings suggest that the deleterious effects of a WD on metabolism are strongly dependent upon the dietary level of isoleucine. Irrespective of sex and strain, mice consuming a WD with a low level of isoleucine are protected from what we normally think of as the inevitable consequences of consuming large quantities of a high-fat, high-sucrose diet, including weight gain and impaired glycemic control. Conversely, a diet with high levels of isoleucine potentiates the negative effects of a WD, with animals gaining more weight and adiposity and becoming more insulin resistant. These effects are broadly in alignment with studies that have shown that increased consumption of all three BCAAs promotes obesity and insulin resistance (Cummings *et al*., 2018; Newgard *et al*., 2009; Solon-Biet et al., 2019), but importantly the negative effects of isoleucine we observed are not coupled to any increase in calorie intake. Indeed, we find that the opposite is true: as levels of dietary isoleucine fall, calorie consumption rises even as metabolic health improves.

This aligns well with the molecular changes we observed, with DBA/2J females having the fewest number of hepatic transcriptional pathways and metabolites altered. Increasing isoleucine levels impaired metabolic health primarily in C57BL/6J mice, where the WD High Ile diet promoted adiposity in males and increased fasting insulin and HOMA2-IR in both sexes. Increased dietary isoleucine altered hepatic transcriptional pathways almost exclusively in C57BL/6J males.

We and others have puzzled over the role of the energy balance hormone FGF21 in the response to dietary protein, with data from C57BL/6J males supporting a model in which induction of FGF21 by PR acts to increase energy expenditure via activation of sympathetic signaling and induction of *Ucp1* in iWAT (Hill et al., 2017; Hill *et al*., 2019). However, we recently suggested that this might be sex and strain-specific, as PR increases blood levels of FGF21 only in C57BL/6J males and not in females or DBA/2J mice (Green *et al*., 2022).

Across all groups, we find that energy expenditure is negatively associated with isoleucine content, and is highest in WD Low Ile-fed mice. However, there is little support for the idea that this increase is mediated by activation of a FGF21-UCP1 axis; dietary levels of isoleucine are negatively correlated with FGF21 levels in C57BL/6J males, but show a U-shaped response curve in DBA/2J males and an inverted U-shaped response curve in females of both strains. FGF21 levels likewise generally do not correlate with *Ucp1* expression, or the expression of other thermogenic and lipolytic genes besides in B6 males. As further evidence that the increased energy expenditure in WD Low Ile-fed mice is not mediated by FGF21, blood levels of FGF21 do not cluster with energy expenditure or amino acid metabolism pathways in our multi-omics analysis; similarly, there is no correlation between circulating FGF21 and dietary Ile levels in humans.

While all protein-containing foods necessarily contain isoleucine, studies of the amino acid methionine have previously suggested that different classes of foods – e.g. plant-derived and animal-derived – may have different amino acid profiles (MacArthur et al., 2021; McCarty et al., 2009). In food item databases we observed that animal items had the highest proportional abundance of isoleucine, although plant-based food items had substantially greater variability in relative isoleucine abundance. Interestingly, investigating human intake data using NHANES led to the observation that lower protein consumers – who as a previous analysis has shown have lower rates of diabetes (Levine *et al*., 2014) – had lower proportional isoleucine intake. Finally, while those in the highest quintile of the HEI had the highest percentage of their diet coming from protein, surprisingly they had the lowest relative consumption of isoleucine. These data are consistent with our findings here and previous studies on isoleucine in humans that support a model in which lower dietary levels of isoleucine promote metabolic health (Deelen et al., 2019; Yu *et al*., 2021).

While drastically reducing dietary levels of isoleucine across the board would require significant changes in the food supply, our analysis within food categories suggests that specific foods in each category may be unusually low in isoleucine content. Similarly, grains and baked goods tended to have lower levels of isoleucine than other food categories. Thus, substituting specific healthy low isoleucine foods may effectively lower dietary levels of isoleucine, and could be a way of rapidly applying these findings to humans. However, such dietary modifications would inevitably alter the intake of many macro- and micro-nutrients, and thus would need to be explicitly tested.

In summary, our results demonstrate that dietary levels of isoleucine strongly regulate the metabolic response to a Western diet across genetic backgrounds and in both male and female mice. Reducing dietary isoleucine broadly reduces adiposity, improves the regulation of blood sugar, and increases energy expenditure. These effects are likely independent of the FGF21-UCP1 axis, as the correlation of FGF21 and *Ucp1* with dietary isoleucine levels is highly dependent upon sex and strain. Exposure to a Western diet containing high isoleucine results in strain-specific detriments in fat mass and insulin resistance. We also find that diet levels of isoleucine as a percentage of protein varies by type of food as well as specific items, and that humans with the highest healthy index scores – diets that most closely align with the Dietary Guidelines for Americans – naturally consume the lowest relative level of dietary isoleucine. Preventing obesity and metabolic syndrome in the face of the cheap high-calorie and high-fat foods ubiquitously available to adolescents is of the highest importance for preventing lifelong obesity and increased risk of many age-related diseases, and our results suggest that reducing dietary isoleucine may be a novel and translatable way to protect adolescents and young adults from the harmful effects of a Western diet on metabolic health.

### Limitations of study

Limitations of our work include the relatively short length of our studies, as well as the use of only two different strains of young mice. Additionally, all of our studies were conducted in the context of a single high-fat, high-sucrose Western diet, and the effects of dietary isoleucine could potentially be different on the background of other unhealthy diets as has been seen for dietary protein (Wali et al., 2021). Additionally, this study did not consider physical activity or age as variables that alter metabolism and dietary responses. Recent data shows that resistance exercise can protect mice from fat accretion observed on a high protein diet (Trautman *et al*., 2023), but how altering isoleucine content would impact the response to exercise, particularly in the context of a Western diet, is unknown. Finally, skeletal muscle retention is vitally important in older adults, and the studies performed here were examined only in young animals.

Our molecular analysis was limited to the liver, and branched-chain amino acids including isoleucine are metabolized in tissues throughout the body (Neinast et al., 2019). Although the phenotypic and molecular correlations we have identified make strong biological sense and align with many of our previous findings, additional work will be required to determine which if any of the findings here are causative. This is true not just with respect to our mouse studies, but also to the human nutrition data analyzed here. Finally, while it appears feasible to construct a diet from natural foods that reduces isoleucine levels, it will require further research to determine if such a diet can improve metabolic health and protect from exposure to an otherwise Western diet.

## Methods

### Animal care, housing and diet

All procedures were performed in conformance with institutional guidelines and were approved by the Institutional Animal Care and Use Committee of the William S. Middleton Memorial Veterans Hospital. Male and female DBA/2J and C57BL/6J mice were procured from the Jackson Laboratory (000664) at six weeks of age. All studies were performed on animals or on tissues collected from animals. All mice were acclimated to the animal research facility for at least one week before entering studies. All animals were housed 2 per cage in static microisolator cages in a specific pathogen-free mouse facility with a 12:12 h light–dark cycle, maintained at approximately 22 °C. At the start of the experiment, mice were randomized to receive the control diet (AA Chow, TD. 140711), 33% isoleucine western diet (WD Low Ile, TD. 170484), 100% isoleucine Western diet (WD Control, TD. 160186), or the 300% isoleucine Western (WD High Ile, TD. 200416) amino acid defined diets; all diets were obtained from Envigo. In the low and high isoleucine diets, nonessential amino acids were adjusted up or down to maintain the same level of dietary nitrogen as the WD Control. Full diet descriptions, compositions and item numbers are provided in **Table 1**.

### In vivo procedures and metabolic phenotyping

Average food consumption was measured over three days in the third week on diet by calculating the difference in food weight between the food put into the cage and that remaining. Food consumption was normalized to weight and lean mass determined at the same time food consumption was measured. Mouse body composition was determined using an EchoMRI Body Composition Analyzer. Glucose and insulin tolerance tests were performed following a 16 hour overnight or 4 hour fast, respectively, and then injecting either glucose (1g/kg) or insulin (0.75U/kg) intraperitoneally (Bellantuono et al., 2020; Yu et al., 2019). Glucose measurements were taken using a Bayer Contour blood glucose meter and test strips. For assays of multiple metabolic parameters (O2, CO2, food consumption, and activity tracking), mice were acclimatized to housing in a Columbus Instruments Oxymax/CLAMS-HC metabolic chamber system for approximately 24 hours, and data from a continuous 24-hour period was then recorded and analyzed.

### Tissue collection for molecular analysis

Terminal submandibular blood was collected on the morning of euthanasia following a 16 hour overnight fast and was then separated into serum. Tissues were rapidly collected and flash frozen in liquid nitrogen, then, with the serum, stored at -80 degrees Celsius until utilized for molecular analysis.

### Assays and kits

Serum insulin was quantified using an ultrasensitive mouse insulin ELISA kit (90080) from Crystal Chem (Elk Grove Village, IL, USA) and blood FGF21 levels were assayed by a mouse/rat FGF21 quantikine ELISA kit (MF2100) from R&D Systems (Minneapolis, MN, USA).

### Quantitative real-time Polymerase Chain Reaction (PCR)

RNA was extracted from iWAT using TRIzol Reagent according to the manufacturer’s protocol (Thermo Fisher Scientific). The concentration and purity of RNA were determined by absorbance at 260/280 nm using Nanodrop (Thermo Fisher Scientific). 1 μg of RNA was used to generate cDNA (Superscript III; Invitrogen, Carlsbad, CA, USA). Oligo dT primers and primers for real-time PCR were obtained from Integrated DNA Technologies (Coralville, IA, USA); sequences are in **<u>Table S2</u>**. Reactions were run on a StepOne Plus machine (Applied Biosystems, Foster City, CA, USA) with SYBR Green PCR Master Mix (Invitrogen). Actin was used to normalize the results from gene-specific reactions.

### Hepatic RNA extraction and –omics analyses

RNA was extracted from liver using TRIzol Reagent according to the manufacturer’s protocol (Thermo Fisher Scientific). The concentration and purity of RNA were determined by absorbance at 260/280 nm using Nanodrop (Thermo Fisher Scientific) and samples were diluted to roughly 1000 ng/µl. RNA was submitted to the University of Wisconsin-Madison Biotechnology Center Gene Expression Center (*RRID:SCR_017757)* & DNA Sequencing Facility *(RRID:SCR_017759).* Each RNA sample was assayed on an Agilent RNA NanoChip to assess its integrity. RNA samples that met Illumina’s TruSeq Stranded Total RNA Reference Guide (#1000000040499 v00) (Illumina, San Diego, CA) quality criteria were prepared for sequencing following the protocol as recommended using a 250ng total RNA input. Libraries were multiplexed and sequenced on the NovaSeq6000 sequencer. Reads were aligned to the mouse (*Mus musculus*) with genome-build GRCm38.p5 of accession NCBI:GCA_000001635.7 and expected counts were generated with ensembl gene IDs (Zerbino et al., 2018).

Analysis of significantly differentially expressed genes (DEGs) was completed in R version 3.4.3 (Team, 2017) using *edgeR* (Robinson et al., 2010) and *limma* (Ritchie et al., 2015). Gene names were converted to gene symbol and Entrez ID formats using the *mygene* package. Genes with too many missing values were removed, if genes were present in less than one diet/strain/sex group they were removed. To reduce the impact of external factors not of biological interest that may affect expression, data was normalized to ensure the expression distributions of each sample are within a similar range. We normalized using the trimmed mean of M-values (TMM), which scales to library size. Heteroscedasticity was accounted for using the voom function, DEGs were identified using an empirical Bayes moderated linear model, and log coefficients and Benjamini-Hochberg (BH) adjusted p-values were generated for each comparison of interest (Benjamini and Hochberg, 2018). DEGs were used to identify enriched pathways, both Gene Ontology (for Biological Processes) and KEGG enriched pathways were determined for each contrast, enriched significantly differentially expressed genes (FDR cutoff=0.1). All genes, log_2_ fold-changes and corresponding unadjusted and Benjamini-Hochberg adjusted p-values can be found in **Tables S3** and **S4**.

Lipids were extracted from liver using a method modified from Matayash et al (2008). Approximately 40mg of liver tissue was added to a bead tube (Qiagen # 13113-50) with 250uL LCMS-grade water and 215uL LCMS-grade MeOH containing 10uL SPLASH lipidomix standard (Avanti #330707) and homogenized in a TissueLyzer (Qiagen) at 4C. 750uL methyl tert-butyl ether was added to each tube, and the tubes were mixed again and allowed to sit on ice for 15 minutes. Then, the samples were centrifuged for 5 minutes at 16100xG at 4C and 450uL of the top organic layer was removed into a new tube. This phase was dried using a speedvac, and lipids were resuspended in 150uL LCMS-grade IPA.

Global lipidomics analysis was performed in positive ion mode on 3uL of lipid extract and in negative ion mode on 5uL of lipid extract. Lipids were separated on a VanGuard BEH C18 precolumn (Waters 18003975) attached to an Acquity BEH C18 column (2.1x100mm, 1.7uM, Waters 186009453) kept at 50C on an Agilent 1290 Infinity II UHPLC system coupled to an Agilent 6545 Q-TOF MS dual AJS ESI mass spectrometer. Mobile phase A consisted of 600:400 Acetonitrile:H2O containing 1mL of formic acid (Fisher A11710X1-AMP) and 0.63g ammonium formate (Honeywell 55674). Mobile phase B consisted of 900:90:10 IPA:Acetonitrile:H2O with 1mL of formic acid and 0.63g ammonium formate. The flow rate was 0.5mL per minute. For both positive and negative mode, the gradient began with 15% mobile phase B, increased to 30% B until 2.4 minutes, and then increased to 48% B until 3 minutes. The gradient increased to 82% B until 13.2 min, and then to 99% B until 13.8 minutes, when it stayed at 99% B until 15.4 minutes, when re-equilibration began and the gradient decreased to 15% B until 20 min.

For both positive and negative ionization mode, the gas temperature was 250C, the flow rate was 12L/min, VCap was 4000V, skimmer was 75V, fragmentor was 190V, and Octapole RF peak was 750V. For positive mode, the sheath gas temperature was 300C, the sheath gas flow rate was 11L/min, and the nebulizer was 35 psig. Reference masses used in positive mode were 121.05 and 922.00 m/z. In negative mode, the sheath gas temperature was 375C, the sheath gas flow rate was 12L/min, and the nebulizer was 30 psig. Reference masses used in negative mode were 112.98 and 966.00 m/z. In both ionization modes, the collision energy for tandem MS was fixed at 25V.

For liver metabolomics, tissue powder was weighed (∼20 mg) on dry ice. The extraction was done by adding 20C methanol:acetonitrile:water (40:40:20) mixture to the powder, followed by vortexing and centrifugation at 16,000 x g for 10 min at 4C. The volume of the extraction solution (mL) was 40 x the weight of tissue (mg) to make an extract of 25 mg tissue per mL solvent. The supernatant (3 mL) was loaded to LC-MS. Samples were analyzed using a quadrupole-orbitrap mass spectrometer (Q Exactive Plus, Thermo Fisher Scientific, San Jose, CA) operating in negative or positive ion modes, coupled to hydrophilic interaction chromatography via electrospray ionization and used to scan from m/z 70 to 1000 at 1 Hz and 140,000 resolution. LC separation was on a XBridge BEH Amide column (2.1 mm x 150 mm, 2.5 mm particle size, 130A° pore size) using a gradient of solvent A (20mM ammonium acetate, 20mM ammonium hydroxide in 95:5 water: acetonitrile, pH 9.45) and solvent B (acetonitrile). Flow rate was 150 ml/min. The LC gradient was: 0 min, 85% B; 2 min, 85% B; 3 min, 80% B; 5 min, 80% B; 6 min, 75% B; 7 min, 75% B; 8 min, 70% B; 9 min, 70% B; 10 min, 50% B; 12 min, 50% B; 13 min, 25% B; 16 min, 25% B; 18 min, 0% B; 23 min, 0% B; 24 min, 85% B; 30 min, 85% B. Autosampler temperature was 4°C.

### Multi-omics analyses

Four data types (metabolomics, transcriptomics, lipidomics, phenotypic outcomes) were obtained from experiments with three factors (strain (B6, DBA), sex, and diet (WD Low Ile, WD Control, and WD High Ile) from 72 mice (6 per diet group of each sex and strain). The data consisted of 534 metabolites, 15636 probes, 344 lipids, and 29 phenotypes.

To identify molecules of interest in each of the -omics datasets, significantly differentially expressed molecules between WD Control and WD High/Low Ile Groups were identified using an empirical Baye’s moderated linear model. The Benjamini-Hochberg method was applied to control false discovery rate, selecting those with adjusted p-value < 0.05 (Benjamini and Hochberg 1995). Transcriptomics, metabolomics and lipidomics data were preprocessed, log2 transformed, z-scale normalized across molecules and samples for each data type individually. Phenotypic outcome data were similarly z-scale normalized just across phenotypes. The data consisted of 5903 inputs across individual mice which had no missing data points including 65 metabolites, 5688 transcripts, 121 lipids, and 29 phenotypes.

To integrate the data, all four datatypes were concatenated for each comparison. Correlations were performed between the 5903 data points using Spearman’s rank (5903 × 5903 = 34,845,409 correlations). Complete hierarchical clustering was used to reorder molecules based on 1 – Spearman correlation between all molecules. The number of clusters were determined by silhouette scores (Rousseeuw, 1987).

All analyses were performed in R (v. 4.2.1) using emmeans (Lenth, 2023) (v. 1.8.4), ComplexHeatmap (Gu et al., 2016) (v. 2.14.0), cluster (Maechler, 2022) (v. 2.1.4). For each cluster, the over representation of KEGG pathways (Kanehisa et al., 2017) from genes were determined using kegga and the gene ontology terms were determined using goana from limma (Ritchie et al., 2015) (v. 3.54.2).

### Transcription factor analysis

Significantly up and downregulated conserved genes were entered into the Mining Algorithm for Genetic Controllers (MAGIC) algorithm (Roopra, 2020). Significant predicted readouts were selected and cross referenced with our hepatic expression analysis to generate a heatmap.

### SHOW study

We analyzed the association between dietary isoleucine intake and FGF21 levels in blood samples collected from 2016−2017 SHOW participants. SHOW is an ongoing population-based health examination survey of non-institutionalized residents of Wisconsin. Detailed survey methods have been previously described (Malecki et al., 2022; Nieto et al., 2010). Survey components relevant to the current analysis included an in-home interview accompanied by measurements of weight and height, and a self-administered questionnaire including the National Cancer Institute’s Diet History Questionnaire, from which specific dietary intake variables were derived.

The study population included 788 individuals who completed all parts of the survey including diet history, blood draw and exam visit. Demographic characteristics of the population have been described previously (Flores et al.). All study protocols were approved by the University of Wisconsin Health Sciences Institutional Review Board, and all participants provided written informed consent as part of the initial home visit. The intake of isoleucine was estimated from the Diet History Questionnaire II (National Cancer Institute) using Diet*Calc software (National Cancer Institute). The estimated levels of isoleucine are expressed as a percentage (%) of the total protein.

FGF21 was quantified from 140 subjects using the previously described ELISA kit and plasma isoleucine determined was determined by the Wisconsin State Laboratory of Hygiene. Multiple linear regression was performed using STATA 17.0 (STATA Corp LLC, College Station, TX, USA) with FGF21 as the outcome and percentage of total protein from isoleucine as the predictor of interest. In the model we also adjusted for age, sex, education, income, total caloric intake and physical activity.

### Diet analyses

For food item analyses, the USDA Food and Nutrient Database SR-28 (NDB) was downloaded. Food items were categorized based on food group codes including dairy and eggs (100), poultry (500), pork (1000), beef (1300), finfish and shellfish (1500), lamb, veal and game (1700), fruits (900), vegetables (1100), nuts and seeds (1200), legumes (1600), cereal grains and pasta (2000). Amino acid analyses in the USDA NDB were performed using three methods for tryptophan, sulfur-containing amino acids and all other amino acids. Tryptophan was measured by alkaline hydrolysis followed by HPLC. Sulfur-containing amino acids were measured by oxidation with performic acid followed by HPLC. All other amino acids were measured by acid hydrolysis followed by HPLC. Values in the USDA NDB are presented as grams of amino acid per 100 grams of food item. Full details on methodology, calculations and data organization are available in the USDA NDB SR28 documentation.

For dietary intake data, data from the National Health and Nutrition Examination Survey (NHANES) was used. Records obtained from the CDC NHANES FTP were read into R using the sasxport.get function from the Hmisc package. Dietary data included two 24-hour dietary recalls conducted by trained dietary interviewers. To assess amino acid composition across dietary records, amino acid data from the USDA NBD SR28 were linked to NHANES food items using SR28 link codes from the Food and Nutrition Database for Dietary Studies (FNDDS). Records were obtained from survey rounds conducted from 2005 to 2012, including four rounds of data release. Individuals who were not pregnant, were over the age of 18 and had two complete 24-hour recalls were included in the analysis, for a total of 23,245 participants. For all nutrient variables including amino acids and total protein, the mean value across two recalls was computed and used for analyses. Healthy Eating Index scores were calculated using the HEI2015_NHANES_FPED function from the dietaryindex R package (v. 1.0.3) using the same set of records described above.

### Statistical analyses

Statistical analysis was conducted using Prism, version 9 (GraphPad Software Inc., San Diego, CA, USA). Tests involving repeated measurements were analyzed with two-way repeated-measures ANOVA, followed by a Tukey–Kramer or Dunnett’s post hoc test as specified. All other comparisons of three or more means were analyzed by one-way ANOVA followed by a Dunnett’s or Tukey–Kramer post hoc test as specified where appropriate. Additional comparisons, if any, were corrected for multiple comparisons using the Bonferroni method. Outliers were determined using GraphPad Prism Grubbs’ calculator (https://www.graphpad.com/quickcalcs/grubbs1/). PCA plots were created using the *“FactoMineR”* package in R. Male and female phenotypic data contained 29 different measures variables and were plotted separately in PCA plots for clarity.

## AUTHOR CONTRIBUTIONS

MET and DWL conceived of and designed the experiments. MET, CLG, MRM, YAH, RB, IJ, MG, EZ, MG, MMS, C-YY, RNM and VF performed the experiments. MET, CLG, MRM, KC, YHA, JAS, IMO, KAM, CJ and DWL analyzed the data. MET, CLG, CJ, JS, IMO, and DWL secured funding and supervised personnel. MET, MM, JAS, IMO, KAM, CJ, and DWL wrote the manuscript.

## ACKNOWLEDGEMENTS

The Lamming lab is supported in part by the NIA (AG056771, AG062328, AG081482, and AG084156), the NIDDK (DK125859), the Wisconsin Partnership Program, and startup funds from UW-Madison. M.E.T. is supported by F99AG083290. CLG was supported in part by Dalio Philanthropies, a Glenn Foundation Postdoctoral Fellowship, and by Hevolution Foundation award HF-AGE AGE-009. RB is supported by F31AG081115. C-YY was supported by T32AG000213 and by F32AG077916. MMS was supported in part by a Supplement to Promote Diversity in Health-Related Research RF1AG056771-06S1. JS is supported by the NIDDK (R01DK133479, P30DK020579), Hatch Grant (WIS04000-1024796); and JDRF (JDRF201309442). JS is a HHMI Freeman Hrabowski Scholar and is an American Federation for Aging Research grant recipient. CJ is supported in part by the NIAAA (R01AA029124). IO was supported by UWCCC Support Grant P30 CA014520 and Wisconsin Head and Neck Cancer SPORE CEP P50DE026787. WAR was supported by U54DK104310 and R01DK131175. Support was provided by the UW-Madison OVCRGE with funding from the Wisconsin Alumni Research Foundation. The authors used the UW-Madison Biotechnology Center Gene Expression Center (RRID:SCR_017757). The Survey of the Health of Wisconsin is funded by the Wisconsin Partnership Program. The Lamming lab was supported in part by the U.S. Department of Veterans Affairs (I01-BX004031), and this work was supported using facilities and resources from the William S. Middleton Memorial Veterans Hospital. The content is solely the responsibility of the authors and does not necessarily represent the official views of the NIH. This work does not represent the views of the Department of Veterans Affairs or the United States Government. Figure 1/panel A Created with BioRender.com released under a Creative Commons Attribution-NonCommercial-NoDerivs 4.0 International license”.

## References

Akter, S., Mizoue, T., Nanri, A., Goto, A., Noda, M., Sawada, N., Yamaji, T., Iwasaki, M., Inoue, M., Tsugane, S., and Japan Public Health Center-based Prospective Study, G. (2021). Low carbohydrate diet and all cause and cause-specific mortality. Clin Nutr 40, 2016–2024. 10.1016/j.clnu.2020.09.022.

Barrington, W.T., Wulfridge, P., Wells, A.E., Rojas, C.M., Howe, S.Y.F., Perry, A., Hua, K., Pellizzon, M.A., Hansen, K.D., Voy, B.H., et al. (2018). Improving Metabolic Health Through Precision Dietetics in Mice. Genetics 208, 399–417. 10.1534/genetics.117.300536.

Batch, B.C., Shah, S.H., Newgard, C.B., Turer, C.B., Haynes, C., Bain, J.R., Muehlbauer, M., Patel, M.J., Stevens, R.D., Appel, L.J., et al. (2013). Branched chain amino acids are novel biomarkers for discrimination of metabolic wellness. Metabolism 62, 961–969. 10.1016/j.metabol.2013.01.007.

Benjamini, Y., and Hochberg, Y. (2018). Controlling the False Discovery Rate: A Practical and Powerful Approach to Multiple Testing. Journal of the Royal Statistical Society: Series B (Methodological) 57, 289–300. 10.1111/j.2517-6161.1995.tb02031.x.

Browning, J.D., Baxter, J., Satapati, S., and Burgess, S.C. (2012). The effect of short-term fasting on liver and skeletal muscle lipid, glucose, and energy metabolism in healthy women and men. J Lipid Res 53, 577–586. 10.1194/jlr.P020867.

Connelly, M.A., Wolak-Dinsmore, J., and Dullaart, R.P.F. (2017). Branched Chain Amino Acids Are Associated with Insulin Resistance Independent of Leptin and Adiponectin in Subjects with Varying Degrees of Glucose Tolerance. Metab Syndr Relat Disord 15, 183–186. 10.1089/met.2016.0145.

Cummings, N.E., Williams, E.M., Kasza, I., Konon, E.N., Schaid, M.D., Schmidt, B.A., Poudel, C., Sherman, D.S., Yu, D., Arriola Apelo, S.I., et al. (2018). Restoration of metabolic health by decreased consumption of branched-chain amino acids. The Journal of physiology 596, 623–645. 10.1113/JP275075.

De Groef, S., Wilms, T., Balmand, S., Calevro, F., and Callaerts, P. (2021). Sexual Dimorphism in Metabolic Responses to Western Diet in Drosophila melanogaster. Biomolecules 12. 10.3390/biom12010033.

Deelen, J., Kettunen, J., Fischer, K., van der Spek, A., Trompet, S., Kastenmuller, G., Boyd, A., Zierer, J., van den Akker, E.B., Ala-Korpela, M., et al. (2019). A metabolic profile of all-cause mortality risk identified in an observational study of 44,168 individuals. Nature communications 10, 3346. 10.1038/s41467-019-11311-9.

Dietary Guidelines for Americans, 2020-2025. (2020). In U.D.o.H.a.H. Services, ed. 9th Edition ed.

Dong, J.Y., Zhang, Z.L., Wang, P.Y., and Qin, L.Q. (2013). Effects of high-protein diets on body weight, glycaemic control, blood lipids and blood pressure in type 2 diabetes: meta-analysis of randomised controlled trials. Br J Nutr 110, 781–789. 10.1017/S0007114513002055.

Felig, P., Marliss, E., and Cahill, G.F., Jr. (1969). Plasma amino acid levels and insulin secretion in obesity. N Engl J Med 281, 811–816. 10.1056/NEJM196910092811503.

Ferraz-Bannitz, R., Beraldo, R.A., Peluso, A.A., Dall, M., Babaei, P., Foglietti, R.C., Martins, L.M., Gomes, P.M., Marchini, J.S., Suen, V.M.M., et al. (2022). Dietary Protein Restriction Improves Metabolic Dysfunction in Patients with Metabolic Syndrome in a Randomized, Controlled Trial. Nutrients 14. 10.3390/nu14132670.

Flippo, K.H., Jensen-Cody, S.O., Claflin, K.E., and Potthoff, M.J. (2020). FGF21 signaling in glutamatergic neurons is required for weight loss associated with dietary protein dilution. Scientific reports 10, 19521. 10.1038/s41598-020-76593-2.

Flores, V., Spicer, A.B., Sonsalla, M.M., Richardson, N.E., Yu, D., Sheridan, G.E., Trautman, M.E., Babygirija, R., Cheng, E.P., Rojas, J.M., et al. (2023). Regulation of metabolic health by dietary histidine in mice. The Journal of physiology 601, 2139–2163. 10.1113/JP283261.

Flurkey, K., Currer, J., and Harrison, D. (2007). The Mouse in Aging Research. In The Mouse in Biomedical Research 2nd Edition, F. JG, ed. (American College Laboratory Animal Medicine), pp. 637–672.

Fontana, L., Cummings, N.E., Arriola Apelo, S.I., Neuman, J.C., Kasza, I., Schmidt, B.A., Cava, E., Spelta, F., Tosti, V., Syed, F.A., et al. (2016). Decreased Consumption of Branched-Chain Amino Acids Improves Metabolic Health. Cell reports 16, 520–530. 10.1016/j.celrep.2016.05.092.

Gannon, M.C., Nuttall, F.Q., Saeed, A., Jordan, K., and Hoover, H. (2003). An increase in dietary protein improves the blood glucose response in persons with type 2 diabetes. Am J Clin Nutr 78, 734–741. 10.1093/ajcn/78.4.734.

Geloneze, B., Vasques, A.C., Stabe, C.F., Pareja, J.C., Rosado, L.E., Queiroz, E.C., Tambascia, M.A., and Investigators, B. (2009). HOMA1-IR and HOMA2-IR indexes in identifying insulin resistance and metabolic syndrome: Brazilian Metabolic Syndrome Study (BRAMS). Arq Bras Endocrinol Metabol 53, 281–287. 10.1590/s0004-27302009000200020.

Green, C.L., Pak, H.H., Richardson, N.E., Flores, V., Yu, D., Tomasiewicz, J.L., Dumas, S.N., Kredell, K., Fan, J.W., Kirsh, C., et al. (2022). Sex and genetic background define the metabolic, physiologic, and molecular response to protein restriction. Cell Metab 34, 209–226 e205. 10.1016/j.cmet.2021.12.018.

Green, C.L., Trautman, M.E., Chaiyakul, K., Jain, R., Alam, Y.H., Babygirija, R., Pak, H.H., Sonsalla, M.M., Calubag, M.F., Yeh, C.Y., et al. (2023). Dietary restriction of isoleucine increases healthspan and lifespan of genetically heterogeneous mice. Cell Metab 35, 1976–1995 e1976. 10.1016/j.cmet.2023.10.005.

Hill, C.M., Laeger, T., Albarado, D.C., McDougal, D.H., Berthoud, H.R., Munzberg, H., and Morrison, C.D. (2017). Low protein-induced increases in FGF21 drive UCP1-dependent metabolic but not thermoregulatory endpoints. Scientific reports 7, 8209. 10.1038/s41598-017-07498-w.

Hill, C.M., Laeger, T., Dehner, M., Albarado, D.C., Clarke, B., Wanders, D., Burke, S.J., Collier, J.J., Qualls-Creekmore, E., Solon-Biet, S.M., et al. (2019). FGF21 Signals Protein Status to the Brain and Adaptively Regulates Food Choice and Metabolism. Cell reports 27, 2934–2947 e2933. 10.1016/j.celrep.2019.05.022.

Huang, J., Liao, L.M., Weinstein, S.J., Sinha, R., Graubard, B.I., and Albanes, D. (2020). Association Between Plant and Animal Protein Intake and Overall and Cause-Specific Mortality. JAMA Intern Med 180, 1173–1184. 10.1001/jamainternmed.2020.2790.

Karusheva, Y., Koessler, T., Strassburger, K., Markgraf, D., Mastrototaro, L., Jelenik, T., Simon, M.C., Pesta, D., Zaharia, O.P., Bodis, K., et al. (2019). Short-term dietary reduction of branched-chain amino acids reduces meal-induced insulin secretion and modifies microbiome composition in type 2 diabetes: a randomized controlled crossover trial. Am J Clin Nutr 110, 1098–1107. 10.1093/ajcn/nqz191.

Kuzuya, M. (2022). Nutritional Management of Sarcopenia and Frailty-Shift from Metabolic Syndrome to Frailty. J Nutr Sci Vitaminol (Tokyo) 68, S67–S69. 10.3177/jnsv.68.S67.

Laeger, T., Henagan, T.M., Albarado, D.C., Redman, L.M., Bray, G.A., Noland, R.C., Munzberg, H., Hutson, S.M., Gettys, T.W., Schwartz, M.W., and Morrison, C.D. (2014). FGF21 is an endocrine signal of protein restriction. J Clin Invest 124, 3913–3922. 10.1172/JCI74915.

Lagiou, P., Sandin, S., Weiderpass, E., Lagiou, A., Mucci, L., Trichopoulos, D., and Adami, H.O. (2007). Low carbohydrate-high protein diet and mortality in a cohort of Swedish women. J Intern Med 261, 366–374. 10.1111/j.1365-2796.2007.01774.x.

Levine, M.E., Suarez, J.A., Brandhorst, S., Balasubramanian, P., Cheng, C.W., Madia, F., Fontana, L., Mirisola, M.G., Guevara-Aguirre, J., Wan, J., et al. (2014). Low protein intake is associated with a major reduction in IGF-1, cancer, and overall mortality in the 65 and younger but not older population. Cell Metab 19, 407–417. 10.1016/j.cmet.2014.02.006.

MacArthur, M.R., Mitchell, S.J., Trevino-Villarreal, J.H., Grondin, Y., Reynolds, J.S., Kip, P., Jung, J., Trocha, K.M., Ozaki, C.K., and Mitchell, J.R. (2021). Total protein, not amino acid composition, differs in plant-based versus omnivorous dietary patterns and determines metabolic health effects in mice. Cell Metab 33, 1808–1819 e1802. 10.1016/j.cmet.2021.06.011.

Maida, A., Zota, A., Sjoberg, K.A., Schumacher, J., Sijmonsma, T.P., Pfenninger, A., Christensen, M.M., Gantert, T., Fuhrmeister, J., Rothermel, U., et al. (2016). A liver stress-endocrine nexus promotes metabolic integrity during dietary protein dilution. J Clin Invest 126, 3263–3278. 10.1172/JCI85946.

Malecki, K.M.C., Nikodemova, M., Schultz, A.A., LeCaire, T.J., Bersch, A.J., Cadmus-Bertram, L., Engelman, C.D., Hagen, E., McCulley, L., Palta, M., et al. (2022). The Survey of the Health of Wisconsin (SHOW) Program: An Infrastructure for Advancing Population Health. Front Public Health 10, 818777. 10.3389/fpubh.2022.818777.

Mancini, M.C., Noland, R.C., Collier, J.J., Burke, S.J., Stadler, K., and Heden, T.D. (2023). Lysosomal glucose sensing and glycophagy in metabolism. Trends in endocrinology and metabolism: TEM 34, 764–777. 10.1016/j.tem.2023.07.008.

Mather, K. (2009). Surrogate measures of insulin resistance: of rats, mice, and men. Am J Physiol Endocrinol Metab 296, E398–399. 10.1152/ajpendo.90889.2008.

McCarty, M.F., Barroso-Aranda, J., and Contreras, F. (2009). The low-methionine content of vegan diets may make methionine restriction feasible as a life extension strategy. Medical hypotheses 72, 125–128. 10.1016/j.mehy.2008.07.044.

Mihaylova, M.M., Chaix, A., Delibegovic, M., Ramsey, J.J., Bass, J., Melkani, G., Singh, R., Chen, Z., Ja, W.W., Shirasu-Hiza, M., et al. (2023). When a calorie is not just a calorie: Diet quality and timing as mediators of metabolism and healthy aging. Cell Metab 35, 1114–1131. 10.1016/j.cmet.2023.06.008.

Mitchell, S.J., Madrigal-Matute, J., Scheibye-Knudsen, M., Fang, E., Aon, M., Gonzalez-Reyes, J.A., Cortassa, S., Kaushik, S., Gonzalez-Freire, M., Patel, B., et al. (2016). Effects of Sex, Strain, and Energy Intake on Hallmarks of Aging in Mice. Cell Metab 23, 1093–1112. 10.1016/j.cmet.2016.05.027.

National Health and Nutrition Examination Survey 2017–March 2020 Prepandemic Data Files Development of Files and Prevalence Estimates for Selected Health Outcomes. (2021). In S. National Center for Health, ed. National Health Statistics Reports. 10.15620/cdc:106273.

Neinast, M.D., Jang, C., Hui, S., Murashige, D.S., Chu, Q., Morscher, R.J., Li, X., Zhan, L., White, E., Anthony, T.G., et al. (2019). Quantitative Analysis of the Whole-Body Metabolic Fate of Branched-Chain Amino Acids. Cell Metab 29, 417–429 e414. 10.1016/j.cmet.2018.10.013.

Newgard, C.B., An, J., Bain, J.R., Muehlbauer, M.J., Stevens, R.D., Lien, L.F., Haqq, A.M., Shah, S.H., Arlotto, M., Slentz, C.A., et al. (2009). A branched-chain amino acid-related metabolic signature that differentiates obese and lean humans and contributes to insulin resistance. Cell Metab 9, 311–326. 10.1016/j.cmet.2009.02.002.

NIDDK (2021). Overweight & Obesity Statistics. https://www.niddk.nih.gov/health-information/health-statistics/overweight-obesity.

Nieto, F.J., Peppard, P.E., Engelman, C.D., McElroy, J.A., Galvao, L.W., Friedman, E.M., Bersch, A.J., and Malecki, K.C. (2010). The Survey of the Health of Wisconsin (SHOW), a novel infrastructure for population health research: rationale and methods. BMC Public Health 10, 785. 10.1186/1471-2458-10-785.

Ramzan, I., Taylor, M., Phillips, B., Wilkinson, D., Smith, K., Hession, K., Idris, I., and Atherton, P. (2020). A Novel Dietary Intervention Reduces Circulatory Branched-Chain Amino Acids by 50%: A Pilot Study of Relevance for Obesity and Diabetes. Nutrients 13. 10.3390/nu13010095.

Ribeiro, R.V., Solon-Biet, S.M., Pulpitel, T., Senior, A.M., Cogger, V.C., Clark, X., O’Sullivan, J., Koay, Y.C., Hirani, V., Blyth, F.M., et al. (2019). Of Older Mice and Men: Branched-Chain Amino Acids and Body Composition. Nutrients 11. 10.3390/nu11081882.

Richardson, N.E., Konon, E.N., Schuster, H.S., Mitchell, A.T., Boyle, C., Rodgers, A.C., Finke, M., Haider, L.R., Yu, D., Flores, V., et al. (2021). Lifelong restriction of dietary branched-chain amino acids has sex-specific benefits for frailty and lifespan in mice. Nat Aging 1, 73–86. 10.1038/s43587-020-00006-2.

Ritchie, M.E., Phipson, B., Wu, D., Hu, Y., Law, C.W., Shi, W., and Smyth, G.K. (2015). limma powers differential expression analyses for RNA-sequencing and microarray studies. Nucleic Acids Research 43, e47–e47. 10.1093/nar/gkv007.

Robinson, M.D., McCarthy, D.J., and Smyth, G.K. (2010). edgeR: a Bioconductor package for differential expression analysis of digital gene expression data. Bioinformatics 26, 139–140. 10.1093/bioinformatics/btp616.

Roopra, A. (2020). MAGIC: A tool for predicting transcription factors and cofactors driving gene sets using ENCODE data. PLoS Comput Biol 16, e1007800. 10.1371/journal.pcbi.1007800.

Roy, S., Sleiman, M.B., Jha, P., Ingels, J.F., Chapman, C.J., McCarty, M.S., Ziebarth, J.D., Hook, M., Sun, A., Zhao, W., et al. (2021). Gene-by-environment modulation of lifespan and weight gain in the murine BXD family. Nat Metab 3, 1217–1227. 10.1038/s42255-021-00449-w.

Seino, Y., Seino, S., Ikeda, M., Matsukura, S., and Imura, H. (1983). Beneficial effects of high protein diet in treatment of mild diabetes. Hum Nutr Appl Nutr 37 *A*, 226-230.

Sluijs, I., Beulens, J.W., van der, A.D., Spijkerman, A.M., Grobbee, D.E., and van der Schouw, Y.T. (2010). Dietary intake of total, animal, and vegetable protein and risk of type 2 diabetes in the European Prospective Investigation into Cancer and Nutrition (EPIC)-NL study. Diabetes Care 33, 43–48. 10.2337/dc09-1321.

Solon-Biet, S.M., Cogger, V.C., Pulpitel, T., Wahl, D., Clark, X., Bagley, E., Gregoriou, G.C., Senior, A.M., Wang, Q.P., Brandon, A.E., et al. (2019). Branched chain amino acids impact health and lifespan indirectly via amino acid balance and appetite control. Nat Metab 1, 532–545. 10.1038/s42255-019-0059-2.

Solon-Biet, S.M., McMahon, A.C., Ballard, J.W., Ruohonen, K., Wu, L.E., Cogger, V.C., Warren, A., Huang, X., Pichaud, N., Melvin, R.G., et al. (2014). The ratio of macronutrients, not caloric intake, dictates cardiometabolic health, aging, and longevity in ad libitum-fed mice. Cell Metab 19, 418–430. 10.1016/j.cmet.2014.02.009.

Solon-Biet, S.M., Mitchell, S.J., Coogan, S.C., Cogger, V.C., Gokarn, R., McMahon, A.C., Raubenheimer, D., de Cabo, R., Simpson, S.J., and Le Couteur, D.G. (2015). Dietary Protein to Carbohydrate Ratio and Caloric Restriction: Comparing Metabolic Outcomes in Mice. Cell reports 11, 1529–1534. 10.1016/j.celrep.2015.05.007.

Team, R.C. (2017). R: A language and environment for statistical computing. R Foundation for Statistical Computing, Vienna, Austria. URL https://www.R-project.org/.

Trautman, M.E., Braucher, L.N., Elliehausen, C., Zhu, W.G., Zelenovskiy, E., Green, M., Sonsalla, M.M., Yeh, C.Y., Hornberger, T.A., Konopka, A.R., and Lamming, D.W. (2023). Resistance exercise protects mice from protein-induced fat accretion. eLife 12. 10.7554/eLife.91007.

Wali, J.A., Milner, A.J., Luk, A.W.S., Pulpitel, T.J., Dodgson, T., Facey, H.J.W., Wahl, D., Kebede, M.A., Senior, A.M., Sullivan, M.A., et al. (2021). Impact of dietary carbohydrate type and protein-carbohydrate interaction on metabolic health. Nat Metab 3, 810–828. 10.1038/s42255-021-00393-9.

White, P.J., Lapworth, A.L., An, J., Wang, L., McGarrah, R.W., Stevens, R.D., Ilkayeva, O., George, T., Muehlbauer, M.J., Bain, J.R., et al. (2016). Branched-chain amino acid restriction in Zucker-fatty rats improves muscle insulin sensitivity by enhancing efficiency of fatty acid oxidation and acyl-glycine export. Molecular metabolism 5, 538–551. 10.1016/j.molmet.2016.04.006.

Yu, D., Richardson, N.E., Green, C.L., Spicer, A.B., Murphy, M.E., Flores, V., Jang, C., Kasza, I., Nikodemova, M., Wakai, M.H., et al. (2021). The adverse metabolic effects of branched-chain amino acids are mediated by isoleucine and valine. Cell Metab 33, 905–922 e906. 10.1016/j.cmet.2021.03.025.

Zerbino, D.R., Achuthan, P., Akanni, W., Amode, M.R., Barrell, D., Bhai, J., Billis, K., Cummins, C., Gall, A., Giron, C.G., et al. (2018). Ensembl 2018. Nucleic Acids Res 46, D754–D761. 10.1093/nar/gkx1098.

Zhu, M., Ji, G., Jin, G., and Yuan, Z. (2009). Different responsiveness to a high-fat/cholesterol diet in two inbred mice and underlying genetic factors: a whole genome microarray analysis. Nutr Metab (Lond) 6, 43. 10.1186/1743-7075-6-43.

