## Supplemental Figures and Table Legends for "Dietary isoleucine content defines the metabolic and molecular response to a Western diet"

### Supplemental Figure 1

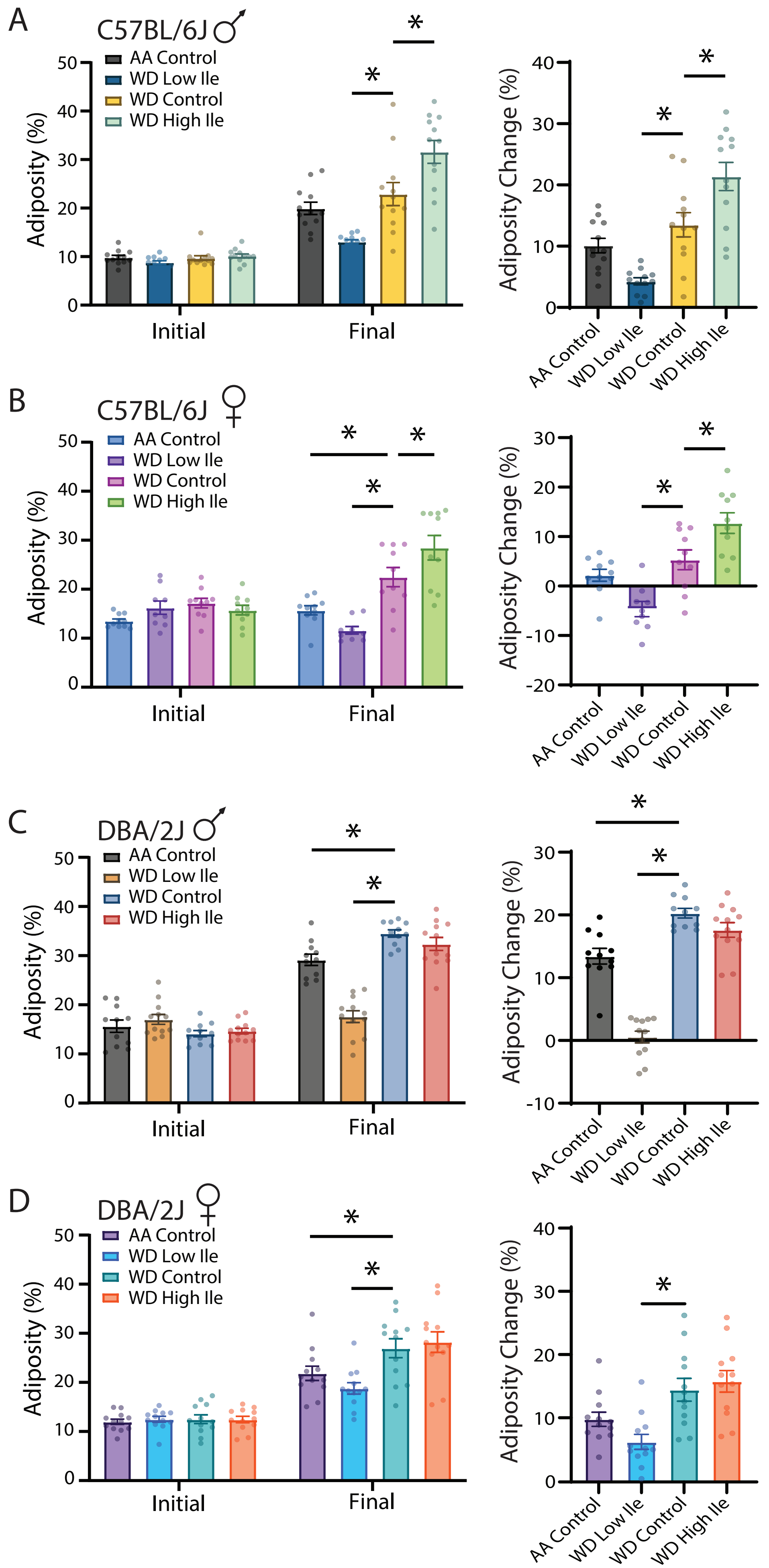

#### Supplemental Figures

**Supplemental Figure 1. Reduced dietary isoleucine promotes leanness in WD-fed mice, while increased dietary isoleucine promotes adiposity in C57BL/6J mice.**

(A-D) Adiposity percent and percent change from week 0 (Initial) to week 12 (Final) in B6 males (A), B6 females (B), DBA males (C), and DBA females (D). n=12 mice per group; $^*p<0.05$, Dunnett's test vs. WD Control post ANOVA, conducted separately for the Final time point (left) and Adiposity Change (%) (right). Data represented as mean ± SEM.

### Supplemental Figure 2

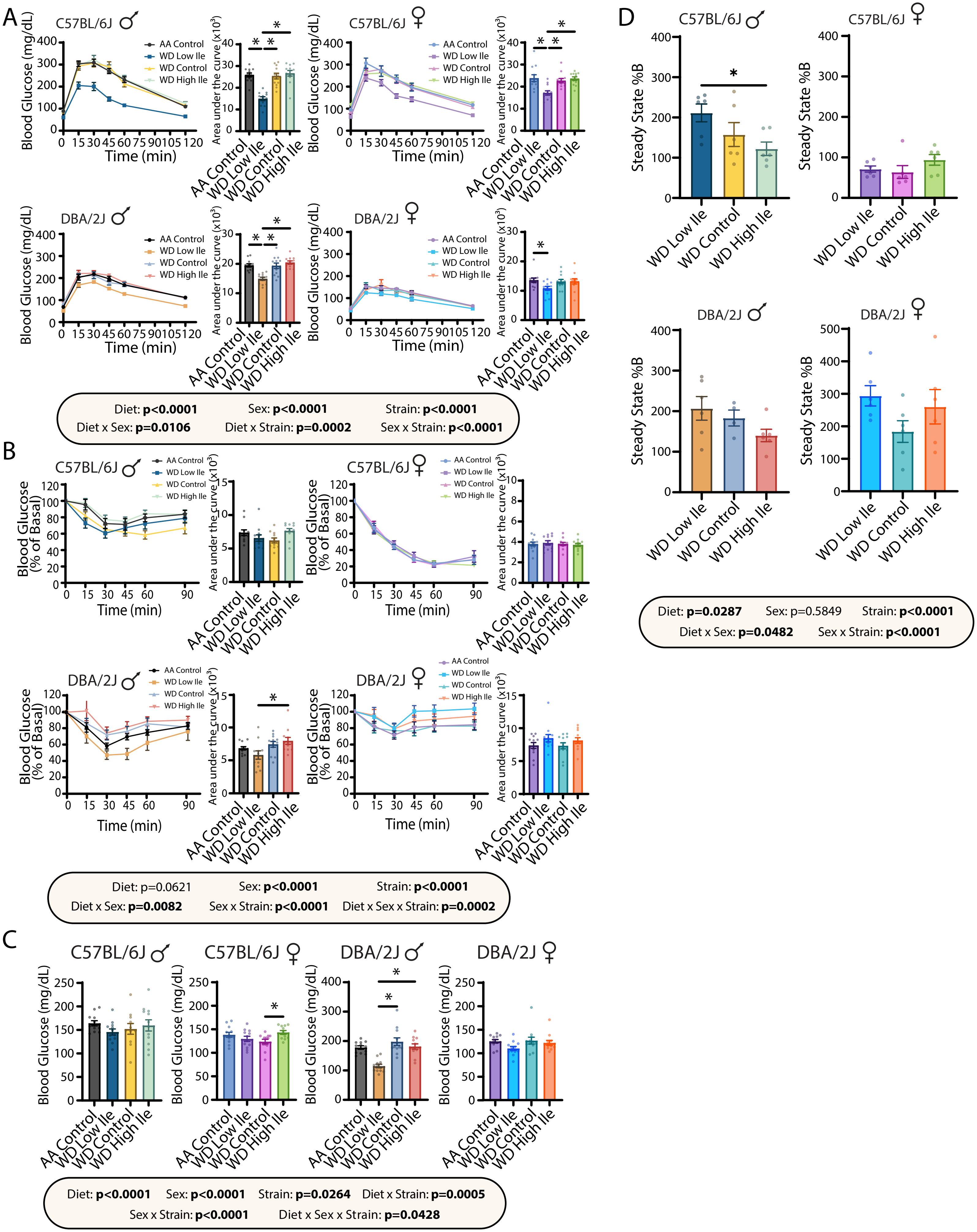

**Supplemental Figure 2. Reduced dietary isoleucine rapidly improves glycemic control.**

(A-B) Glucose (A) and insulin (B) tolerance tests in mice fed the indicated diets for 3 weeks and 4 weeks, respectively, with quantified area under the curve (AUC). (C) 4 hour fasting blood glucose levels. (D) Steady state %B calculated from HOMA2-IR parameters calculated after 12 weeks on the indicated diets from the values indicated in **Figure 2**. (A-D) n=5-12 mice per group. Statistics for the overall effects of diet, sex, and strain represent the p value from a three-way ANOVA, $*p<0.05$, Tukey's test post ANOVA for each sex/strain group shown. Data represented as mean ± SEM.

### Supplemental Figure 3

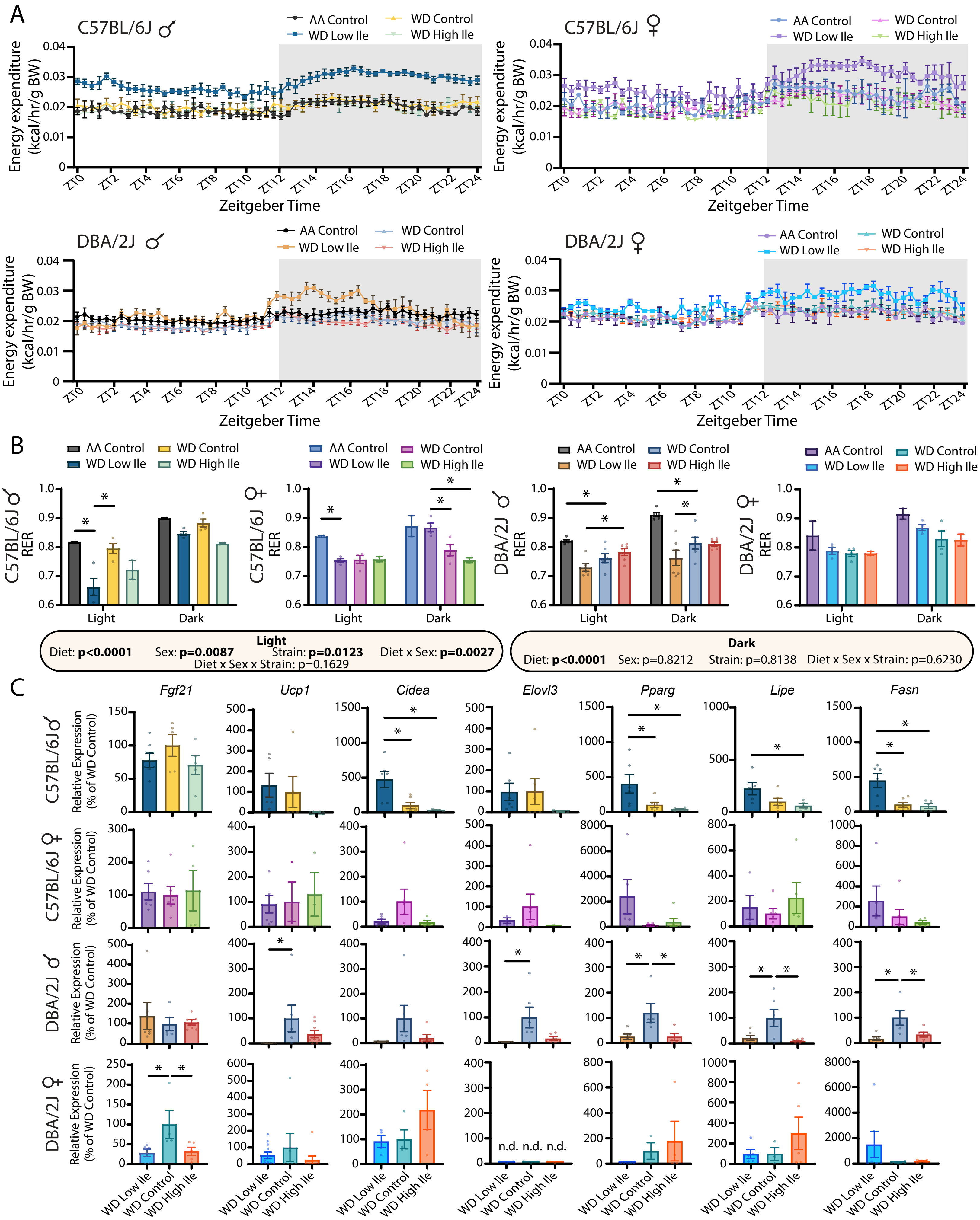

**Supplemental Figure 3. Dietary levels of isoleucine impact energy balance.**

(A) 24-hour metabolic chamber analysis of energy expenditure per body weight. (B) Light and dark average respiratory exchange ratio values. (C) Inguinal white adipose tissue gene expression. n=2-6 mice per group. (B) Statistics for the overall effects of diet, sex, and strain represent the p value from a three-way ANOVA; *$p<0.05$, Tukey's test post ANOVA for each light/dark cycle separately. (C) *$p<0.05$, Tukey's test post ANOVA. Data represented as mean ± SEM.

### Supplemental Figure 4

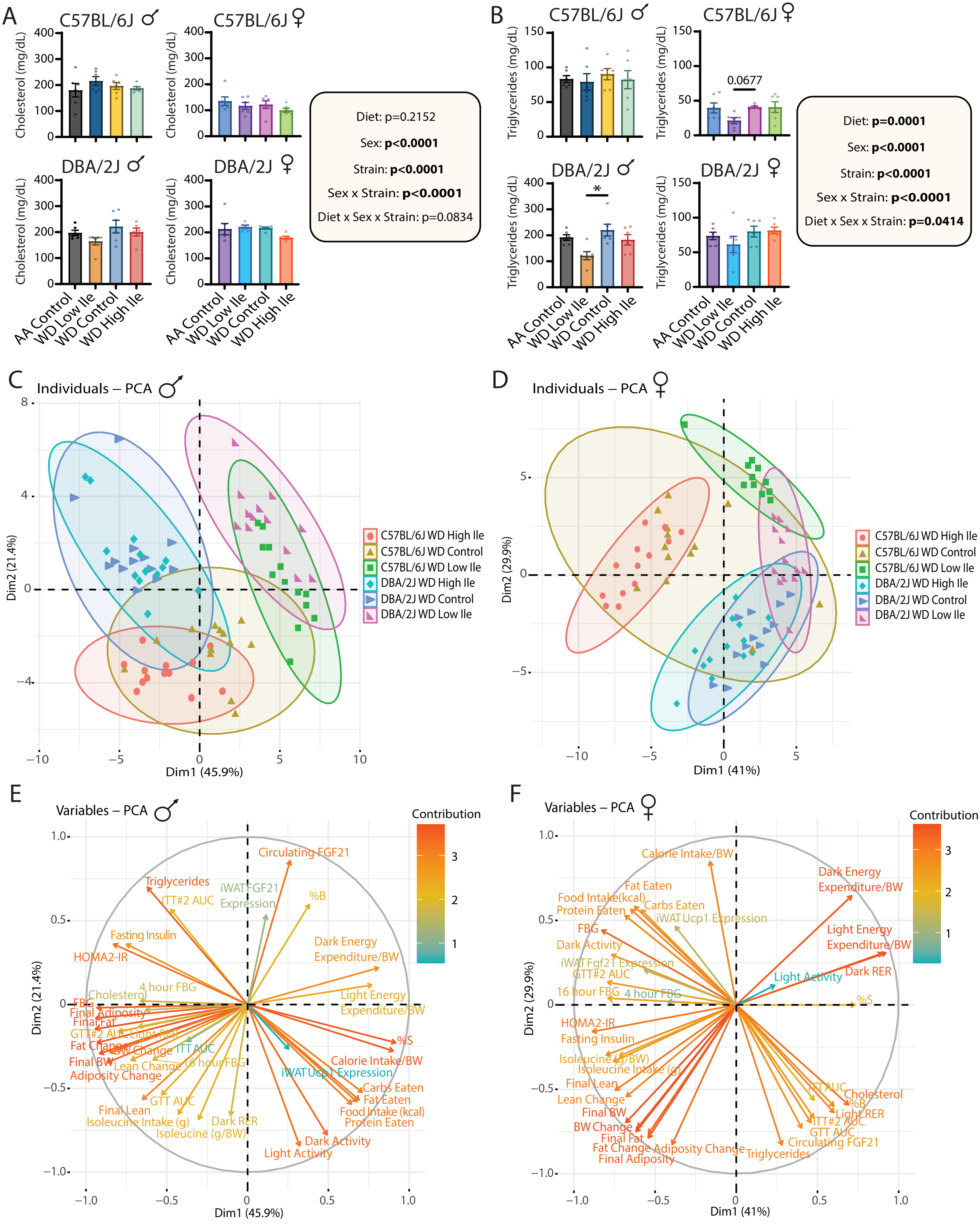

**Supplemental Figure 4. Principle component analysis of the effects of dietary isoleucine level, and the impact of dietary isoleucine level on circulating lipids, in different sexes and strains.**

(A-B) Circulating fasting (A) cholesterol and (B) triglycerides. (C-D) Principle component analysis of phenotypes in (C) males and (D) females, and (E-F) the variables contributing to the PCA spread. (A-B) n=6 mice per group. (C-F) n=2-12 mice per phenotype/group. (A-B) Statistics for the overall effects of diet, sex, and strain represent the p value from a three-way ANOVA; $^*p<0.05$, Tukey's test post ANOVA for each sex/strain group shown. Data represented as mean ± SEM.

### Supplemental Figure 5

A

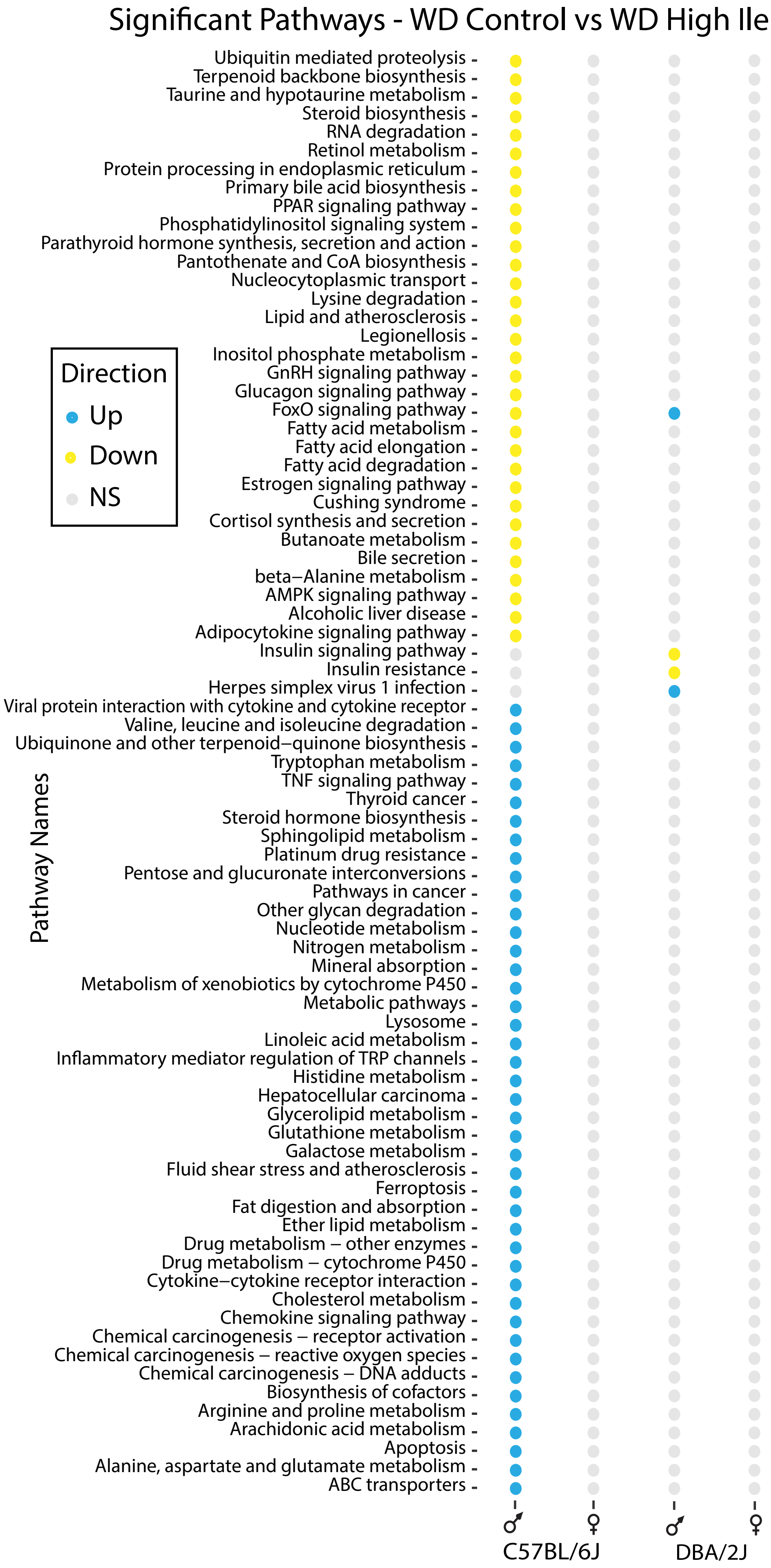


B

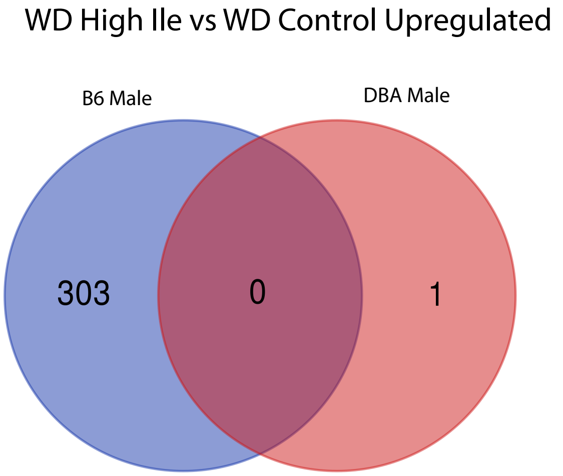


C

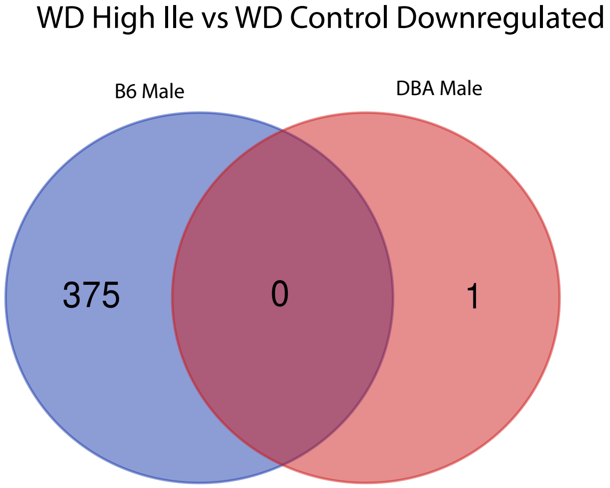

**Supplemental Figure 5. Increased dietary isoleucine has sex and strain specific effects on hepatic gene expression.**

(A) KEGG pathway analysis of significantly altered genes in WD High Ile-fed mice relative to WD Control-fed mice in each sex and strain. (B) Shared (B) upregulated and (C) downregulated genes in B6 and DBA males.

### Supplemental Figure 6

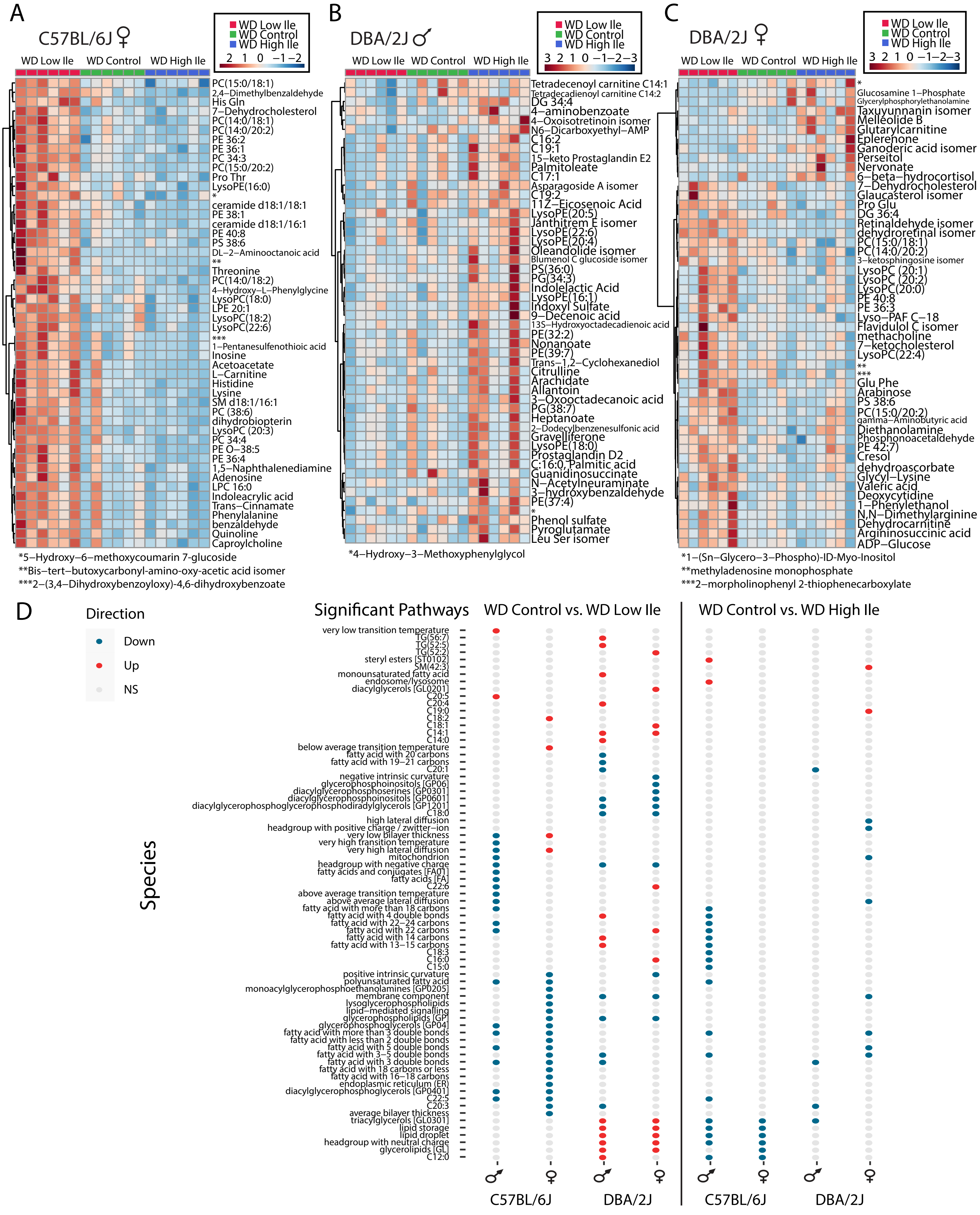

**Supplemental Figure 6. Metabolites and lipids altered by isoleucine.**

(A-C) Top 50 hepatic metabolites altered in response to dietary isoleucine in B6 females (A), B6 males (B), and DBA females (C). (D) Pathways identified as significantly altered in at least one group based on lipidomics analysis of mice of each sex and strain on the indicated diets.

### Supplemental Figure 7

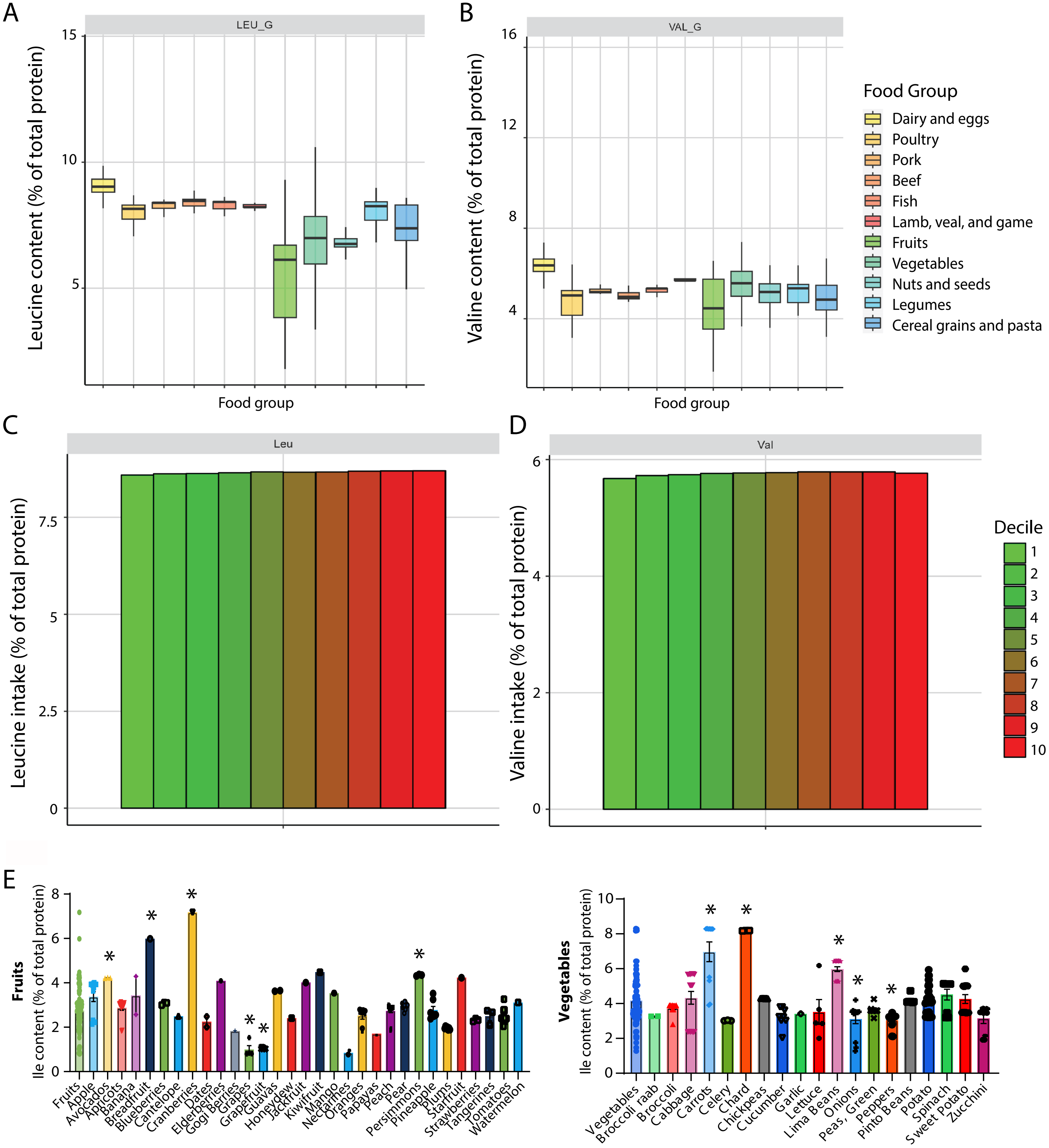

**Supplemental Figure 7. National Health and Nutrition Examination Survey (NHANES) Diet analyses.**

(A-B) Leucine (A) and valine (B) as a percentage of total protein across food groups. (C-D) Relative leucine (C) and valine content (D) per decile of protein intake from NHANES dietary data. (E) Relative isoleucine content in commonly consumed fruits and vegetables, *$p<0.05$, Dunnett's post-test vs. Fruits or Vegetables.

**Supplemental Table Legends**

Table S1: Diet composition.

Table S2: Oligo dT primers and primers for RT-qPCR displayed in Supplemental Figure 3C.

Table S3: Upregulated transcripts ($log_2$ fold change).

Table S4: Downregulated transcripts ($log_2$ fold change).

Table S5: KEGG Enriched Pathways used to generate Figure 4A, Supplemental Figure 5A.

Table S6: Top 50 Metabolites for each sex and strain used to generate Figure 4B, Supplemental Figure 6A-C.

Table S7: Conserved genes, metabolites, and lipids used to generate Figure 5G-H.

Table S8: Transcripts, metabolites, and lipids used to generate Venn Diagrams in Figure 5A-F.

Table S9: Genes, metabolites, lipids, and phenotypes present in each megacluster in Figure 6A.

Table S10: RT-qPCR p-values from 3-way ANOVA displayed in Supplemental Figure 3C.
